# Improving acetate metabolism of *P. putida* KT2440 by evolutionary and rational engineering

**DOI:** 10.64898/2026.08.21.746131

**Authors:** Melanie Filbig, Luisa Wachtendonk, Luca Hampe, Isabel Bator, Josefin Johnsen, Elsayed T. Mohamed, Nicolás Gurdo, Johannes Parschau, Pablo I. Nikel, Adam M. Feist, Till Tiso, Lars M. Blank

## Abstract

Acetate is a promising carbon source for microbial biotechnology as it can be produced sustainably from lignocellulosic biomass or C1 gases. Since acetate is directly activated to acetyl-CoA, it is especially suitable for producing acetyl-CoA-derived products, showcased here with the production of 3-(3-hydroxyalkanoyloxy) alkanoic acids (HAAs). *P. putida* KT2440 can natively metabolize acetate, but the weak acid has also inhibitory effects on microbial growth. We present an in-depth study on the physiology of *P. putida* KT2440 using acetate as carbon and energy source and evaluate acetate as feedstock for the biosynthesis of HAAs.

Initially, a rational engineering approach to overexpress acetyl-CoA synthetase for acetate activation resulted in an improved growth rate of 16% and reduced lag phase by six hours. To further increase the performance of *P. putida* KT2440 on acetate, adaptive laboratory evolution was performed. This resulted in an improvement in the growth rate from 0.4 h^-1^ to 0.6 h^-1^ and enabled growth on up to 12.5 g L^-1^ acetate with a shortened lag phase compared to the wild type. Whole-genome sequencing revealed mutations in proteins involved in gene expression regulation and signal transduction. This evolutionary engineering approach informed the deletions of *gacS* and *crc*, which resulted in a reduction in the lag phase from seven hours to one hour and an improvement of the growth rate by 25 %, matching the growth properties of the evolved clones. Using the evolved strains for the production of HAAs resulted in faster biomass and product formation with product titers reaching up to 94 % of that of the wild type. In conclusion, we identified mechanisms in the acetate metabolism of *P. putida* KT2440 and improved the growth performance of the strain by rational and evolutionary engineering, demonstrating the potential of the promising, but challenging 3^rd^ generation feedstock acetate.

## 1 Introduction

Although glucose is a common biotechnological substrate it is discussed controversially in the food vs. fuel debate and its high price in Europe during 2023-2024 (Vesper 2025) further motivates the search for alternative substrates. The substrate choice significantly impacts bioprocess costs, as it can account for up to 70 % of the total expenses (Hepner 1996). The chosen substrate also influences other parameters, including microbial growth and, thus, space-time yield, as well as product and biomass yields, and therefore the sustainability of the process. Special attention is currently paid to substrates derived from biomass, *i.e*., lignocellulose, although the only wheat straw-based bioethanol plant in Europe closed in late 2023 (Clariant 2023). Alternatively, C1-gasses can be used, such as those from the exhaust gas generated during steel production, as demonstrated by LanzaTech’s bioethanol production process (Köpke and Simpson 2020). C1 and C2 substrates such as methane, methanol, formic acid are also reinvestigated, as are ethylene glycol, ethanol, and acetate. Acetate (CH_3_COO^-^) with alternative chemical and microbial synthesis routes might have a price advantage (300 – 450 $ per ton (Kiefer et al. 2021)) over glucose at ≈700 $ per ton (Chemanalyst), or at least has a lower spread in price fluctuation. Although 85 % of the total acetate production relies on the carbonylation of fossil-derived methanol (Kalck et al. 2020), several sustainable chemical and biological alternatives exist, including the hydrolysis of lignocellulosic biomass (Mills et al. 2009, Gong et al. 2016), anaerobic digestion (Braun 2013), and syngas fermentation (Kantzow et al. 2015). Furthermore, acetate production from CO_2_ by microbial electrosynthesis is possible (Gildemyn et al. 2015). Current research also focusses on establishing routes to oxidize CH_4_ from fossil resources and emissions to acetate using methanotrophs (Soo et al. 2016, Cai et al. 2019). The use of a carbon source, which can be generated from several waste streams, *i.e.*, C1-gasses or agricultural waste containing lignocellulose, can contribute to the overall sustainability and carbon-balance of the biotechnological process. Besides biological acetate generation, sustainable chemical catalysis can also be used, including the conversion of CO_2_ and CH_4_ present in biogas to acetic acid (Martín-Espejo et al. 2022), the generation of acetic acid from CO_2_ by electrochemical reduction (Hori et al. 2002), or the production of acetate from CO_2_-derived methyl-formate (Jürling-Will et al. 2022). To reduce the price even further, coal to acetate is reemerging in some markets, opening possibilities for acetate-based biotechnology, however, without the benefit of defossilization.

Microbial growth on acetate involves three key steps: (I) acetate entering the cell, (II) activation into acetyl-CoA, and (III) assimilation *via* the glyoxylate and TCA cycles. Acetate enters the cell either by diffusion (undissociated form) or active transport using H+/monocarboxylate symporters (PMCT) or acetate permeases (ActP). Once inside, acetate is activated into acetyl-CoA through two ATP-dependent pathways: the acetyl-CoA synthase has high affinity for acetate and requires two ATP per acetyl-CoA (Wolfe 2005, Kiefer et al. 2021), or the AckA-PTA pathway, in which acetate is first converted to acetyl-phosphate, and second converted to acetyl-CoA (Kutscha and Pflügl 2020). Acetyl-CoA then enters the TCA cycle and glyoxylate shunt, enabling the formation of C4 molecules from C2 substrates (Kiefer et al. 2021). These C4 molecules are then used for amino acid biosynthesis and gluconeogenesis, necessary for growth on C2 substrates.

Many biotechnologically relevant organisms, such as *Escherichia coli*, *Saccharomyces cerevisiae*, *Ustilago maydis*, *Corynebacterium glutamicum*, and *Pseudomonas putida*, can use acetate as their sole carbon and energy source. Acetate has been used to produce various value-added compounds including succinic acid (Li et al. 2016), glycolic acid (Li et al. 2019), isopropanol (Yang et al. 2020), isobutanol (Song et al. 2018), amino acids (Jo et al. 2019), malic acid (Kövilein et al. 2021), lipids (Zhang et al. 2019), and polyhydroxyalkanoates (PHAs) in organisms like *E. coli*, *A. oryzae*, and *R. glutinis*. An *acs*-overexpression mutant of *P. putida* KT2440 was applied to produce medium chain-length polyhydroxyalkanoates (mcl-PHA) using sodium acetate as sole carbon source (Yang et al. 2019). Furthermore, Arnold et al. (2019) used *P. putida* KT2440 to produce rhamnolipids from acetate and other small organic acids. However, the toxicity of acetate, which naturally occurs as a preservative in the food industry, at high concentrations limits its use as carbon source, necessitating hosts with higher tolerance and efficient metabolization. Overexpressing genes for acetate activation into acetyl-CoA (Lin et al. 2006, Lee et al. 2018, Yang et al. 2019) or enhancing the glyoxylate shunt can improve acetate utilization. For example, *E. coli* with overexpressed glyoxylate shunt genes increased itaconate production from acetate (Noh et al. 2018). However, reducing fluxes of gluconeogenic reactions was shown to have no impact on acetate conversion to acetone (Yang et al. 2019).

3-(3-Hydroxyalkanoyloxy) alkanoic acids (HAAs) are esters of two β-hydroxy fatty acids (Abdel-Mawgoud et al. 2010). Besides their surface-active properties, rendering them biosurfactants, the molecule is a promising precursor for chemical conversion due to the several chemical groups contained. By converting HAAs chemocatalytically, bioplastic, biokerosene, and bio-hybrid fuel molecules can be synthesized (Beydoun and Klankermayer 2019, Meyers et al. 2019, Mensah et al. 2020, Tiso et al. 2021). Naturally, HAAs occur in the biosynthesis pathway of the well-studied biosurfactants rhamnolipids. Since the best-studied rhamnolipid producer *P. aeruginosa* PAO1 is an opportunistic human pathogen, recombinant production of HAAs and rhamnolipids was established in *P. putida* KT2440 by heterologous expression of either the acyltransferase *rhlA* only or combined expression with rhamnosyltransferase I *rhlB* (Ochsner et al. 1995, Wittgens et al. 2011, Blesken et al. 2020, Tiso et al. 2020). By expressing *rhlA* under the control of three stacked synthetic promoters in the *att*Tn7-site of *P. putida* KT2440, up to 940 mg L^-1^ HAAs were produced from 10 g L^-1^ glucose (Blesken et al. 2020). Since *P. putida* KT2440 is natively able to use acetate as sole carbon- and energy source, the aim of this work was to improve the growth and production performance on acetate. First, biomass and product synthesis were investigated and compared to results obtained on glucose. Two different approaches were then used with the aim of improving the growth behavior of *P. putida* KT2440 on acetate. The first approach focused on rational engineering, while the second included adaptive laboratory evolution (ALE), followed by whole genome sequencing and reverse engineering. Since the effective equimolar conversion of acetate to acetyl-CoA by one or two steps makes acetate a suitable substrate for the production of acetyl-CoA-derived bioproducts, the production of HAAs from acetate was showcased.

## 2 Material and methods

### 2.1 Bacterial strains, media, and growth conditions

All strains used in this work are listed in Table 1. Strains of *E. coli* and *P. putida* KT2440 were routinely cultivated in lysogeny broth (LB), according to Lennox (1955). For the preparation of solid medium, 1.5 % agar was added prior to autoclaving. While *P. putida* was cultivated at 30°C, *E. coli* was cultivated at 37°C. Antibiotics were added if required to avoid plasmid loss and for selective purposes in the following concentrations: 50 µg mL^-1^ kanamycin, 30 µg mL^-1^ gentamycin, and 100 µg mL^-1^ ampicillin.

**Table 1:**
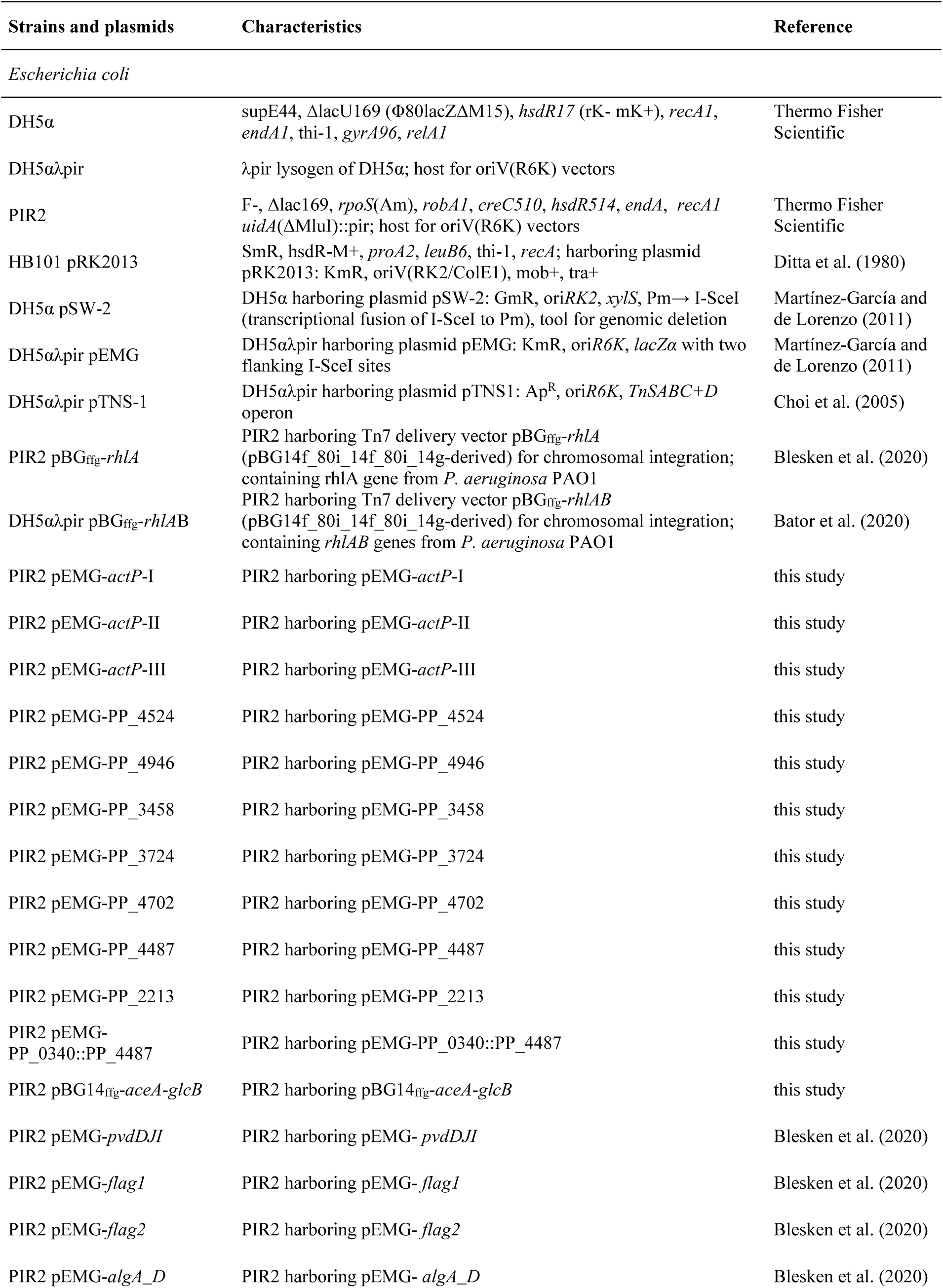
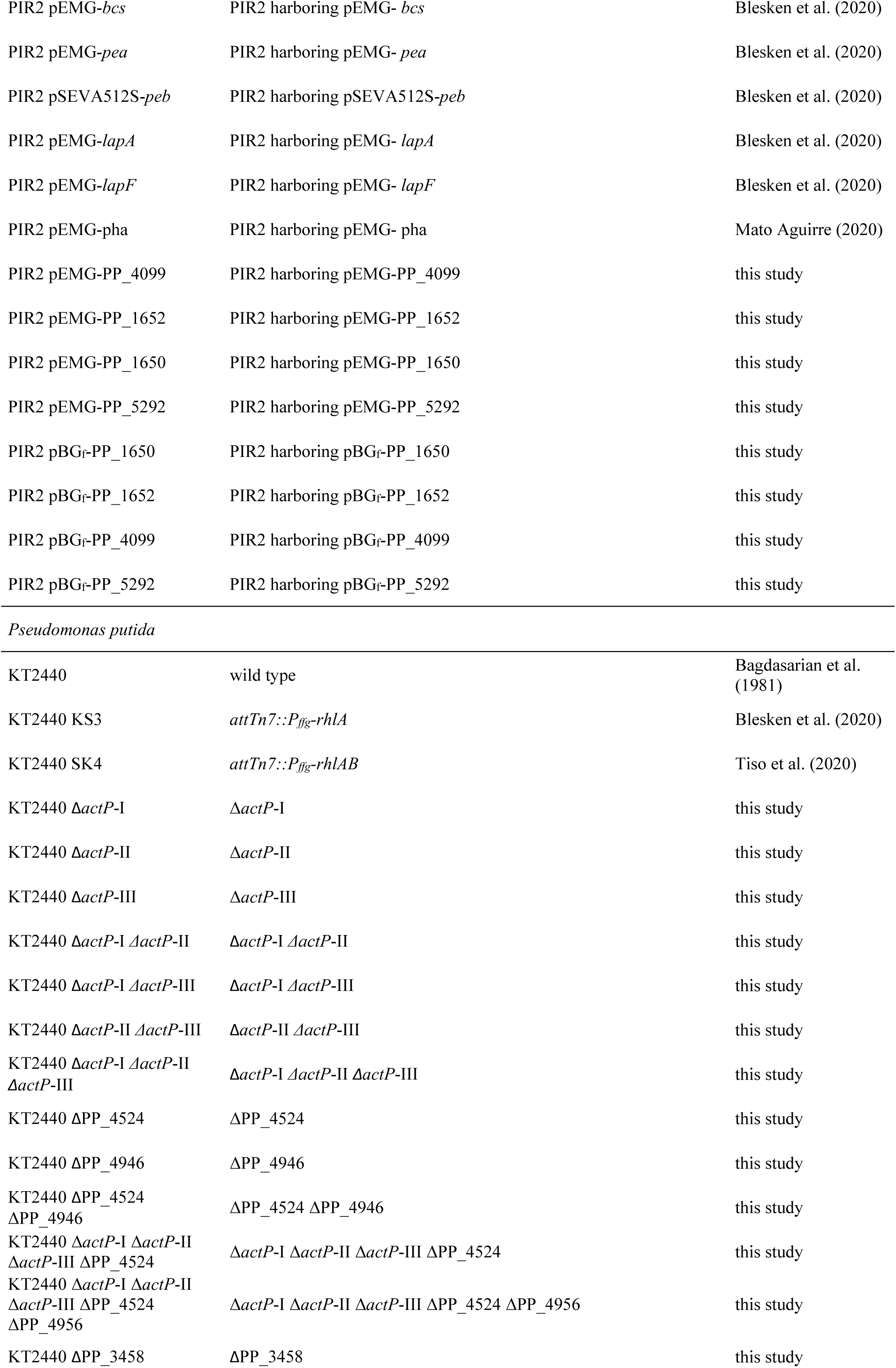
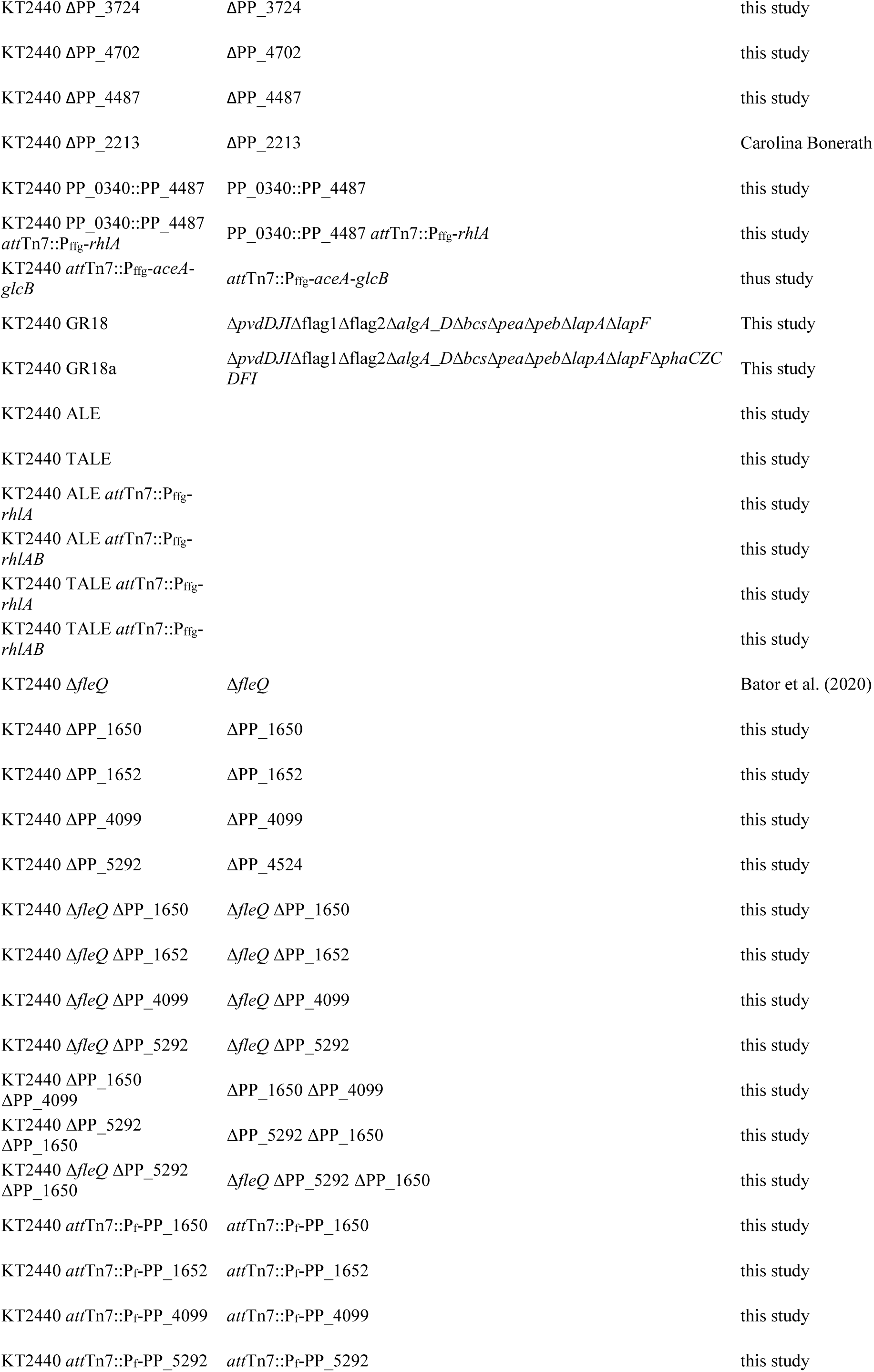
Bacterial strains used in this work.

For the selection of *P. putida* strains after mating procedures, cetrimide agar (Sigma-Aldrich, St. Louis, MO, USA) was used to avoid growth of *E. coli*. Growth and production experiments were performed using mineral salts medium (MSM) with a final composition (per L) of 1.55 g K_2_HPO_4_, 0.85 g NaH_2_PO_4_·2H_2_O, 2.0 g (NH_4_)_2_SO_4_, 0.1g MgCl_2_·6H_2_O, 10 mg EDTA, 2 mg ZnSO_4_·7H_2_O, 1 mg CaCl_2_·2H_2_O, 5 mg FeSO_4_·7H_2_O, 0.2 mg Na_2_MoO_4_·2H_2_O, 0.2 mg CuSO_4_·5H_2_O, 0.4 mg CoCl_2_·6H_2_O, and 1 mg MnCI_2_·2H_2_O (Hartmans et al. 1989), and 10 g glucose or 5 g acetate for pre-cultures or different concentrations of glucose and acetate for main cultures as indicated. All shaken cultures were inoculated to an optical density (OD_600_) of 0.1 to start cultivation.

Liquid cultivations were performed in 100 mL or 250 mL shake flasks for pre-cultures and 500 mL shake flasks for main cultivations, each with 10 % filling volume at 250 or 300 rpm and a shaking diameter of 50 mm. For small-scale cultivations, 24-deep well plates (SystemDuetz; Enzyscreen B.V., Heemstede, The Netherlands) with 1.5 mL filling volume were cultivated at 300 rpm and a shaking diameter of 50 mm. Online growth monitoring was performed using a Growth Profiler 960 (Enzyscreen B.V., Heemstede, The Netherlands) in white 24-deep well plates with clear bottom with 1.5 mL filling volume at 225 rpm and a shaking diameter of 50 mm. For determination of culture turbidity, *i.e*., biomass formation, green value (GV) was extracted from images of the culture every 30 minutes by the image analysis software “Growthviewer” (Enzyscreen B.V., Heemstede, The Netherlands).

Production of metabolic CO_2_ during cultivation was monitored using BCP-CO_2_ sensors and the BlueVIS software (BlueSens, Herten, Germany). Cultivation was performed in 1 L shake flasks filled with 50 ml medium containing 0.17 Cmol of the respective carbon source at 150 rpm with a shaking diameter of 50 mm.

Online monitoring of oxygen and carbon dioxide transfer rates in shaking flasks was performed using Kuhner TOM (Adolf Kühner AG, Birsfelden, Switzerland). The filling volume and shaking frequency were set as indicated.

### 2.2 Plasmid and strain construction

For construction of plasmids, which was done using NEBuilder® HiFi DNA Assembly Kit (New England Biolabs, Ipswich, MA, USA), the NEBuilder Assembly online tool was used. DNA fragments were amplified using Q5 High-Fidelity DNA Polymerase (New England Biolabs, Ipswich, MA, USA) according to the manufacturer’s instructions. Primers were ordered as custom DNA oligonucleotides (Eurofins Genomics, Ebersberg, Germany) and are listed in supplementary information, Table S 1. Following DNA assembly, the products were introduced into chemically competent *E. coli* PIR cells *via* heat shock, adhering to the protocol of Hanahan (Green and Sambrook 2018). Verification of positive clones was performed by colony PCR using OneTaq® 2X Master Mix with Standard Buffer (New England Biolabs, Ipswich, MA, USA). Cell material from colonies underwent lysis with alkaline polyethylene glycol before PCR according to Chomczynski and Rymaszewski (2006).

For the creation of gene deletions, the I-*Sce*I-based system (Martínez-García and de Lorenzo, 2011) was employed. 500 bp regions located upstream and downstream of the target deletion region, referred to as TS1 and TS2 regions, were amplified from *P. putida* KT2440 genomic DNA (isolated using the Monarch® Genomic DNA Purification Kit, New England Biolabs, Ipswich, MA, USA) and cloned into the suicide delivery vector pEMG, which has previously been linearized *via* restriction digest with EcoRI (New England Biolabs, Ipswich, MA, USA). The conjugation protocol from Wynands et al. (2018) was used for the transfer of resulting pEMG deletion plasmids from *E. coli* PIR2 into the relevant *Pseudomonas*. Kanamycin-sensitive clones underwent the removal of the pSW-2 plasmid through multiple transfers of cells into fresh LB medium without antibiotics, followed by re-verification using colony PCR. The system was further used for genomic integration into PP_0340 (Köbbing et al. 2024). Therefore, the DNA sequence to be integrated was amplified *via* PCR and cloned in between TS1 and TS2.

Rhamnolipid and HAA producers were generated by integrating *rhlAB* into the *att*Tn7-site downstream of *glmS* using the mini-Tn7 delivery transposon vector pBG14f_80i_14f_80i_14g (Köbbing et al. 2020), as described by Bator et al. (2020). To construct the mini-Tn7 delivery transposon vector for the integration of *rhlA*, the acyltransferase responsible for the formation of HAA, pBGffg (Köbbing et al. 2020) was linearized *via* PCR using primers MF222 and MF223. *rhlA* was amplified from *P. aeruginosa* PAO1 using primers MF224 and MF225. The plasmid pSK02 or pKS03, housing the mini-Tn7 delivery transposon, was integrated into the genome of *P. putida* KT2440 mutants through transposition. The identification of rhamnolipid- or HAA-producing colonies was carried out as described by Bator et al. (2020). The colonies exhibiting the highest product titers were selected after cultivation in LB medium containing 10 g L^-1^ glucose. Genomic integration into *att*Tn7 was also used to overexpress the glyoxylate shunt by integrating a second copy of the genes *aceA* and *glcB* using the before mentioned delivery transposon vector.

### 2.3 Adaptive laboratory evolution

Adaptive laboratory evolution (ALE) and tolerance adaptive laboratory evolution (TALE), which were used to improve the performance of *P. putida* KT2440 on acetate, were performed using an automated liquid handler platform (LaCroix et al. 2015, Mohamed et al. 2017, Mohamed et al. 2019). Cultivation was conducted as a batch fermentation at 30°C at a stirring speed of 1200 rpm. Cells were transferred to a new flask when they reached a cell density of around 0.47, corresponding to OD_600_ = 2 on a common 1 mm pathlength benchtop reader. Four populations of each, *P. putida* KT2440, *P. putida* KT2440 GR18, and *P. putida* KT2440 GR18a, were evolved for 612-638, 528-637, and 507-546 generations, respectively (Table 2). Two different ALE experiments were performed. In the first experiment, performed to improve the growth rate of *P. putida* KT2440 on acetate, a constant acetate concentration of 0.083 M was applied. In a second experiment, performed to improve the tolerance of *P. putida* KT2440 to higher acetate concentrations, the acetate concentration in the medium was increased whenever growth rate was restored to WT growth rate, starting from 0.083 M acetate. An aliquot of 2% of the cell culture was transferred into fresh medium when a threshold of OD_600_=0.86 was reached. Periodically, glycerol stocks were made to preserve the progress, monitor the development of the mutations, and for genome sequencing after 24 transfers (int1) and after 109 transfers (end) for the ALE experiments and after 34 (int1), 64 (int2) and 108 (end) transfers for the TALE experiment.

**Table 2:**
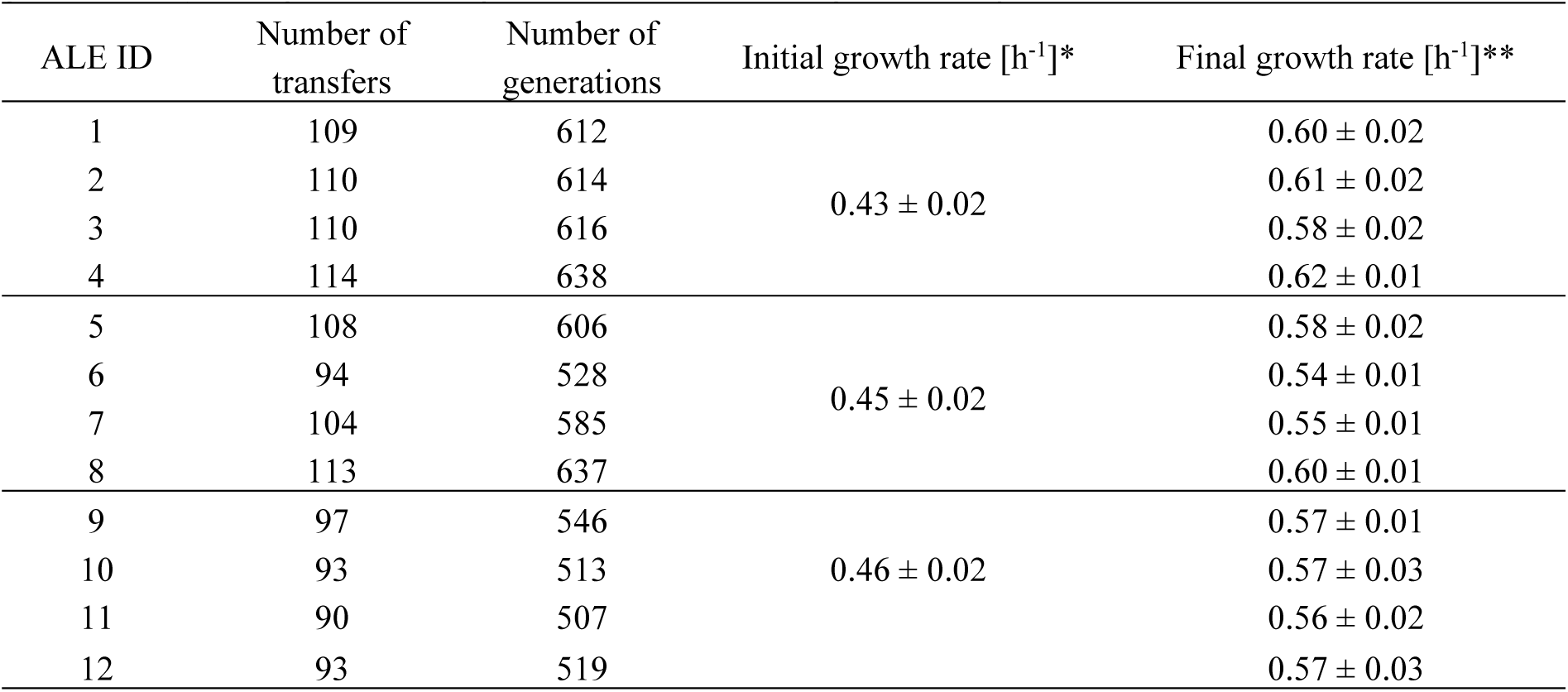
Growth rates of each individually evolved lineage of *P. putida* KT2440 (ALE ID 1-4), *P. putida* KT2440 GR18 (ALE ID 5-8), and *P. putida* KT2440 GR18a (ALE ID 9-12) ALE experiments, including the number of flasks and generations. *) Average of the initial growth rate for all four lineages. **Average of the five last flasks.

After the end of the ALE experiments, ten single colonies were isolated from each frozen sample and investigated regarding their phenotype. One single colony of each isolate was used for genome sequencing.

### 2.4 Whole genome sequencing

Genomic DNA was isolated using Monarch® Genomic DNA Purification Kit (New England Biolabs, Ipswich, MA, USA) prior to assessment of the DNA quality by determination of A260/280 using a Nanodrop (Thermo Fisher scientific, Waltham, USA). After generation of paired-end sequencing library generation using the Plexwell 384 Library Prep kit (SeqWell), genome sequencing of ALE isolates was performed using the NextSeq high Output kit V2 a 300 cycle (150 bp x 2) from Illumina (San Diego, USA) to identify mutations responsible for phenotypic changes of *P. putida* KT2440 after ALE on acetate. Reads resulting from sequencing were compared to the genome sequence of *P. putida* KT2440 (NC-002947.4) using an inhouse pipeline (Phaneuf et al. 2019) to find deletions, insertions (InDels), and single nucleotide polymorphisms (SNP). The mutation tables can be found on the ALEdb database (ALEdb.org) under project name “Putida_Acetate”.

### 2.5 Gene expression studies

RNA sequencing was performed to identify genes involved in the metabolism of acetate. Samples for RNA sequencing were taken during cultivation in the early exponential phase when the culture had an OD of 1.0 - 1.5. The corresponding cell count was determined using the BactoBox® (SBT Instruments, Copenhagen, Denmark). 1 ml sample was pelleted by centrifugation at 13,300 rpm for 2 min, and the supernatant was discarded prior to snap-freezing in liquid nitrogen. Samples were stored at -80°C until sent for RNA isolation, rRNA depletion and RNA sequencing, which was performed by Eurofins Genomics Germany GmbH (Ebersberg, Germany), as well as gene expression analysis.

### 2.6 Analytical methods

Biomass formation was monitored during shake flask cultivations by measuring optical density at 600 nm (OD_600_). A correlation between OD_600_ and cell dry weight (CDW) was determined with *P. putida* KT2440 on glucose to a value of 0.356 g_CDW_/OD unit.

#### 2.6.1 Analysis of acetate

Acetate concentration in the supernatant was analyzed using HPLC-RI. Samples from cultivation were taken and centrifuged at 13,3000 rpm for 3 min prior to filtration in glass vials using ROTILABO® Cellulose acetate (CA) syringe filters with 0.2 µm pore size (Carl Roth GmbH + Co. KG, Karlsuhe, Germany). Samples were analyzed in an HPLC consisting of a pump module, autosampler, column oven, UV detector and an RefractoMax 521 RI detector (all Thermo Fisher Scientific, Waltham, MA, USA) with a Metab-AAC 300 x 7.8 mm column (particle size: 10 µm, ISERA GmbH, Düren, Germany). Elution was performed with 5 mM H_2_SO_4_ at a flow rate of 0.6 mL min^-1^ at 40°C.

#### 2.6.2 Analysis of rhamnolipids and HAAs

HAA and mono-RL concentrations were determined using reversed phase HPLC-CAD, similar to the method described by Behrens et al. (2016). For sample preparation, cultivation broth was mixed 1:1 with acetonitrile and stored at 4°C overnight before it was centrifuged at 13,300 rpm for 3 min and filtered into glass vials with Phenex RC syringe filters (0.2 µm, Ø 4 mm) (Phenomenex, Torrance, CA, USA). An Ultimate3000 HPLC system consisting of a dual gradient pump, an autosampler and a column compartment connected to a Corona Veo Charged Aerosol Detector (all Thermo Fisher Scientific, Waltham, MA, USA) was used. For chromatographic separation, a NUCLEODUR C18 Gravity 150 x 3 mm column (particle size: 3 µm, Macherey-Nagel GmbH & Co. KG, Düren, Germany) was used. The flow rate of the analytical gradient and the inverse gradient was set to 0.425 mL min^-1^; the column oven temperature was set to 40°C. 0.2 % (v/v) formic acid in acetonitrile (A) and 0.2% (v/v) formic acid in ultra-pure water (B) were used for gradient elution. The gradient was set to 70 % A and 30 % B at the start. The share of A linearly increased to 80 % in the first eight minutes and remained constant for 1 min, before increasing to 100 % until min 13. After three minutes, A decreased back to 70 % in 1.5 min and remained constant until the measurement ended after 25 min. The injection volume was set to 3 µl.

To equalize the altering solvent concentration in the CAD and guarantee a constant solvent share of 85 %, an inverse gradient was applied, which was calculated by the software in the mode ‘keep solvent composition’ (Chromeleon 7.2.10). The offset was determined experimentally to 778 µl. The share of A in the inverse gradient was 100 % in the beginning and decreased to 90 % from 1.83 min to 9.83 min. After 10.83 min, the share of A decreased further to 70 %, before it increased back to 100 % after 19.33 min. 1-monoolein was used to generate a calibration function.

## 3 Results

### 3.1 Physiological effects of acetate on *P. putida* KT2440

Acetate is known for its toxicity to microbial cells due to different mechanisms, even at low concentrations (Pinhal et al. 2019). To investigate, which acetate concentrations can be tolerated by *P. putida* KT2440, the bacterium was cultivated on different concentrations of acetate as single carbon source and in combination with glucose as second substrate (Figure 1b, c).

**Figure 1:**
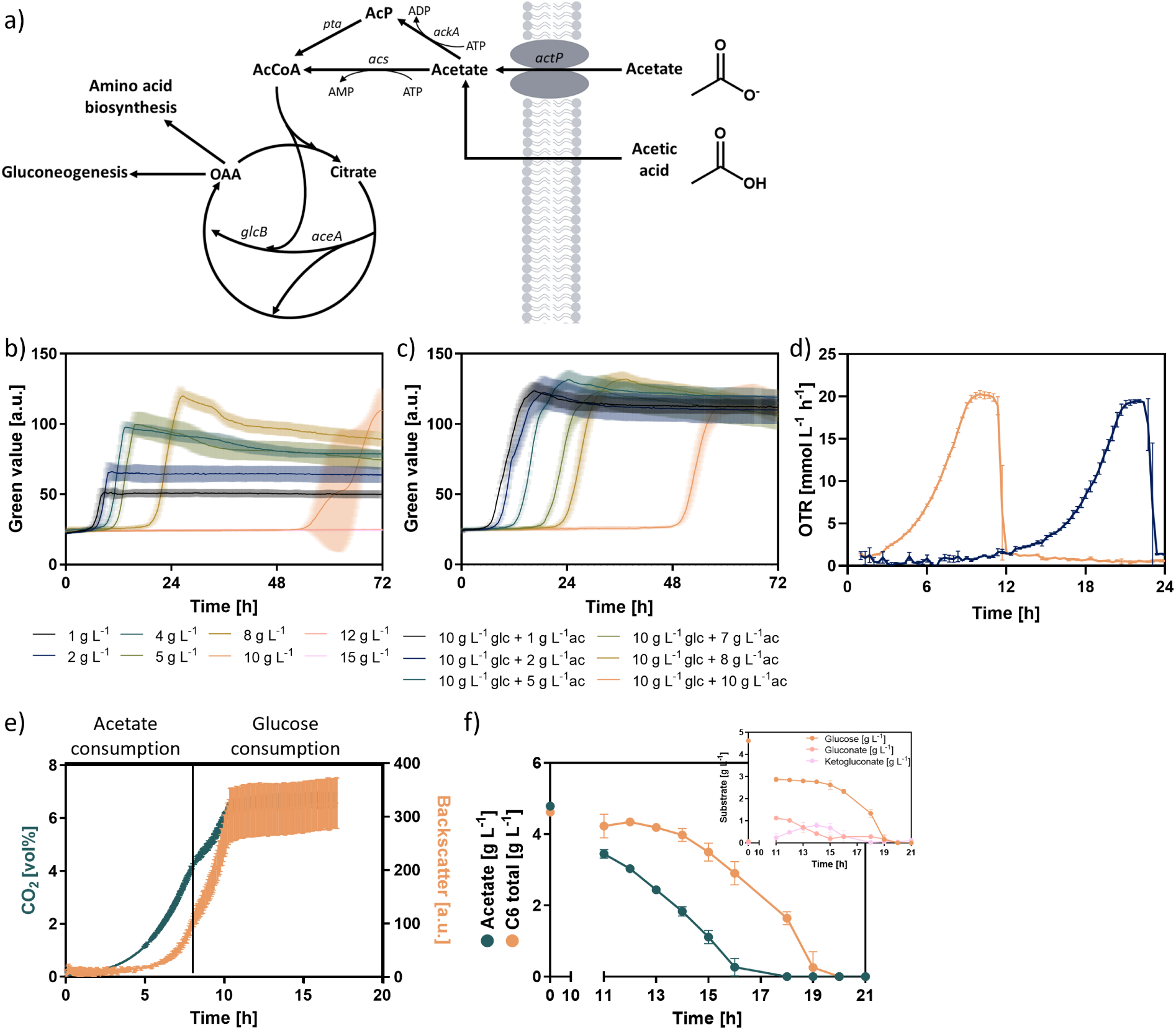
Acetate metabolism and physiological characterization of *P. putida* KT2440 on acetate as single or co-substrate. a) schematic representation of acetate metabolism. While the uptake of acetate requires transporters as acetate permease ActP, the undissociated form, acetic acid, can freely enter the cell *via* diffusion. Two possibilities exist for acetate activation: (I) *via* acetyl-CoA synthetase (*acs*) or (II) *via* acetate kinase (*ackA*) and phosphate acetyltransferase (*pta)*. Acetyl-CoA is assimilated *via* the glyoxylate shunt, yielding oxaloacetate (OAA). *actP* = acetate permease, *acs* = acetyl-CoA synthetase, *ackA* = acetate kinase, *pta* = phosphate acetyltransferase, *aceA* = isocitrate lyase, *glcB* = malate synthase G, AcP = acetyl-phosphate, AcCoA = acetyl-CoA, OAA = oxaloacetate. b)-c) Growth of *P. putida* KT2440 on different acetate concentrations with b) Acetate as sole carbon source and c) as co-substrate to 10 g L^-1^ glucose. d) OTR measurement of a *P. putida* KT2440 cultivation on 5 g L^-1^ acetate inoculated from a pre-culture grown on glucose (blue) or on acetate (orange). e)-f) Cultivation of *P. putida* KT2440 KS3 on equimolar amounts of glucose and acetate with e) online measurement of CO_2_ (blue) and backscatter (orange) during growth on 2.1 g L^-1^ glucose and 2.1 g L^-1^ acetate. CO_2_ was measured using BCP-CO_2_ sensors (BlueSens gas sensor GmbH, Herten, Germany), backscatter was determined using the Cell Growth Quantifier (Scientific Bioprocessing, Inc., Baesweiler, Germany). Cultivation was performed in 1 L shake flasks with 5 % filling volume at 150 rpm. The vertical line indicates the transition between the two growth phases. b) Concentrations of acetate, glucose, gluconate, and 2-ketogluconate during growth on 5 g L^-1^ glucose and 5 g L^-1^ acetate. Cultivation was performed in 500 ml shake flasks with 10 % filling volume. Error bars represent standard deviations of two biological replicates

Acetate affects the growth of *P. putida* KT2440 as a sole or second substrate. When acetate is used as sole carbon source, the lag phase prolongs with increasing acetate concentration. The growth rate of 0.33 h^-1^ was unaffected at a concentration up to 5 g L^-1^ but was reduced to 0.29 h^-1^ at 8 g L^-1^ and 0.23 h^-1^ at 10 g L^-1^. Biomass concentration increased with increasing acetate concentration, indicating that acetate can be metabolized and used for biomass production by the strain and that a higher substrate availability correlates with higher biomass formation.

While on a concentration of 10 g L^-1^ acetate, biomass formation starts after more than 48 hours, no biomass formation was observed on 12 g L^-1^ and 15 g L^-1^ acetate within 72 hours. When acetate is used in combination with glucose, an inhibitory effect is also visible: the higher the applied acetate concentration, the longer the lag phase. Also, the growth rate is affected even at a concentration of 5 g L^-1^ acetate. While the cells exhibit a growth rate of 0.31 h^-1^ at 2 g L^-1^ added acetate, the growth rate drops to 0.26 h^-1^ at 5 g L^-1^ and to 0.22 h^-1^ at 7 g L^-1^. Although it is difficult to say for sure, a potential diauxie in the growth profile at 9, 11, and 18 h in the curves for 1 g L^-1^, 2 g L^-1^, and 5 g L^-1^ added acetate, respectively, might hint at a sequential use of the two available carbon sources glucose and acetate. Due to the nature of the used device, no statement can be made at this point about the biomass concentration achieved as the maximal detectable green value was reached with the lowest acetate concentration, already. All in all, the data show that biomass formation is possible on acetate, but that acetate negatively affects growth of *P. putida* KT2440 in a concentration-dependent way.

In order to reduce the lag phase on acetate, pre-culture management on acetate was investigated. An initial approach for cultivation in minimal medium was a first pre-culture in complex medium from a frozen stock and this served as staring material for a second pre-culture in glucose-containing minimal medium. Thus, cells were physiologically adapted to the minimal medium already when the main cultivation was inoculated, but not to the C2-substrate. Therefore, a third pre-culture in acetate-containing minimal medium, inoculated from the second pre-culture, was considered to allow for accustoming of *P. putida* KT2440 to the organic acid. To investigate the effect of this third pre-culture, *P. putida* KT2440 cells from a pre-culture on glucose or a pre-culture on acetate were used as inoculum for a cultivation on 5 g L^-1^ acetate with online measurement of O_2_ and CO_2_. While the strain only exhibited a lag phase of around 1.6 hours when inoculated from a pre-culture on acetate, the strain required about 9 hours to start growth on acetate when inoculated from a pre-culture on glucose, extending the cultivation time from 12 h to 24 h (Figure 1d). Maximal OTRs reached under both conditions were comparable but showed oxygen limitation, independent of the pre-culture management. Thus, once the cells start growing, the cell’s metabolism is not influenced by the different pre-culture strategies. These results motivated the use of a third pre-culture in minimal medium with acetate as sole carbon source to allow physiological adaptation to the C2-compound prior to main cultivations and, thus, to reduce the cultivation time. Furthermore, these results show that short-term adaptation to acetate is possible and can improve the performance. Skipping the rich media or glucose containing pre-cultures was not explored.

### 3.2 Revealing growth characteristics of *P. putida* KT2440 during co-utilization of acetate and glucose

Given evidence of a sequential metabolization of glucose and acetate (Figure 1c), the potential diauxie was further investigated. To do this, the amount of CO_2_ released during cultivation on a mixture of acetate and glucose was monitored as an indicator of metabolic activity with parallel measurement of biomass formation. In this experiment, a mutant of *P. putida* capable of the production of 3-(3-hydroxyalkanoyloxy) alkanoic acids (HAA), *P. putida* KT2440 KS3, was used. On the mixture of acetate and glucose, the strain exhibited a noticeable diauxic growth (Figure 1ea). Monitoring glucose and acetate concentrations in the medium during a parallel cultivation with sampling revealed that acetate is consumed in the first growth stage. Glucose, however, is oxidized to gluconate and 2-ketogluconate, which is the natural fate of the hexose in the periplasm of *P. putida* KT2440 (del Castillo et al. 2007), but not further metabolized in the presence of acetate. When acetate is depleted, gluconate and 2-ketogluconate are consumed by the strain to continue growth in the second growth phase (Figure 1f). Since organic acids are preferred carbon sources over glucose for *Pseudomonas* (Rojo 2010), this result is not surprising.

In the growth phase on acetate, the strain reached a backscatter value of 100 while 4.3 vol% CO_2_ are produced. In the second growth phase, the strain reaches a final density of 320, while the CO_2_ concentration increases to 6.5 vol%. In total, an increase in the density of 220 and an increase in the CO_2_ signal of 2.2 % resulted from the second growth phase. The results thus indicate that biomass formation from acetate is less efficient than from glucose since more biomass was generated in the second growth phase. Furthermore, the data revealed that CO_2_ formation is almost twice as much on acetate (4.3 vol%) than on glucose (2.2 vol%), indicating an altered metabolism on the C2 compound. Key CO_2_-delivering reactions in the cell, catalyzed by isocitrate dehydrogenase and α-ketoglutarate dehydrogenase in the TCA cycle, often result in carbon loss during energy production. The anaplerotic reaction on acetate is the glyoxylate shunt, which can bypass the CO_2_-delivering steps, but it exhibits also reduced energy (NADH) generation. Therefore, some flux through the lower TCA cycle is necessary to ensure enough NADH, despite carbon loss.

These results are in accordance with wet-lab and dry-lab data from Ziegler et al. (2023), showing a higher CO_2_ generation of *P. putida* KT2440 KS3 from acetate. In this previous study, altered fluxes through the TCA cycle on acetate, resulting in elevated CO_2_ formation, were shown. In a shake flask experiment on 5 g L^-1^ acetate or glucose, a lower biomass yield on acetate was confirmed: while a biomass yield of 0.41 g_CDW_ g_substrate_^-1^ was determined on glucose, a yield of 0.27 g_CDW_ g_substrate_^-1^ was determined on acetate (data not shown), confirming the described trend. A reason for the lower biomass formation on the C2 compound might be its lower energy content on one side (38 ATP for 1 glucose vs. 10 ATP for 1 acetate (Seong et al. 2020)) and the higher energy demand for acetate metabolization on the other side. This might also explain a higher flux through the NADH-generating but CO_2_-releasing reactions mentioned above, catalyzed by isocitrate dehydrogenase and α-ketoglutarate dehydrogenase. Besides exhibiting toxic effects, including lower growth rates and higher lag phases as determined in section 3.1, the C2 compound thus also leads to a less efficient carbon-to-biomass conversion.

In summary, it was revealed that *P. putida* KT2440 KS3 shows diauxic growth on a mixture of acetate and glucose, preferring acetate. Further, biomass formation from acetate was found to be less efficient than from glucose, mainly caused by an elevated CO_2_ production to meet the higher energy demands. These findings motivate a detailed consideration and optimization of the acetate metabolism with the aim to improve growth performance, including biomass yield.

### 3.3 Acetate impairs biomass and HAA production in *P. putida* KT2440 KS3

Intracellular acetate is directly converted into acetyl-CoA, rendering it an optimal substrate for acetyl-CoA derived products. The formation of the acetyl-CoA derived product HAA was demonstrated already: when comparing the performance of *P. putida* KT2440 KS3 on acetate to the conventional carbon source glucose in a shake flask experiment, Ziegler et al. (2023) determined reduced biomass yields, growth rates, as well as yields of the product HAA. Further, HAA production from acetate lacks far behind theoretical computations: while on glucose, the theoretical HAA yield was calculated to be 0.68 Cmol Cmol^-1^, the theoretical product yield was reduced to 0.59 Cmol Cmol^-1^ on acetate (Ziegler et al. 2023). Reasons for a reduced HAA formation from acetate are similar to the reasons for a reduced biomass formation given in section 3.2.

Investigating HAA formation on different acetate-to-glucose ratios in wet-lab experiments also revealed a negative impact of acetate on the biomass titers, as well as the specific HAA yields (Figure 2). When glucose was reduced and acetate increased to 50 % each of the total carbon amount, the biomass dropped to 71 % while the specific HAA yield even dropped to 59 % of the yield reached on glucose, which is a yield reduction of 82 %, although the overall available carbon was kept constant.

**Figure 2:**
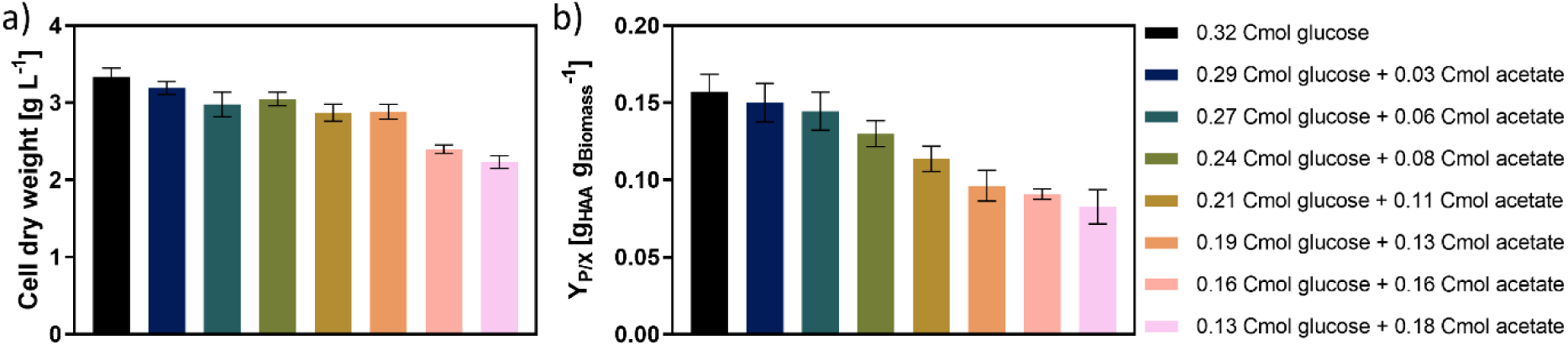
Biomass and product formation of *P. putida* KT2440 KS3 on different combinations of glucose and acetate as substrates. a) Cell dry weight and b) specific HAA yield. Error bars represent standard deviation of n=3.

Overall, acetate strongly impacted the performance of *P. putida* KT2440 as a biocatalyst. Even the presence of small amounts of acetate as co-substrate negatively impacted obtained biomass and product yields. Several reasons explaining these findings might exist. On the one hand, the toxic effect of the protonated form of acetic acid, as acidification of the cytoplasm, might play a role, albeit not a major one. Furthermore, the lower energy content of acetate, with a higher energy demand for metabolizing the C2-compound, might explain the reduced yields regarding biomass and HAA. The altered metabolism also plays an important role: when more of the available carbon is “burnt” for CO_2_ generation in favor of NADH generation, less carbon is available for forming biomass or products. The combination of these effects probably explains the low yields observed here.

### 3.4 <u>Ac</u>etate <u>t</u>ransport <u>p</u>roteins (ActP) unlikely to be involved in acetate uptake

To have the possibility to improve the acetate metabolism of *P. putida* towards faster assimilation and conversion, the fate of acetate in the cell was investigated. The metabolic pathway of acetate consumption was considered step by step, using deletion mutans to identify bottlenecks and overexpression to address them. The first step in metabolizing acetate is the transfer across the cell membrane. In *P. putida* KT2440, three genes are predicted to encode acetate permeases, facilitating acetate diffusion across the membrane, by sequence comparison: *actP*-I (PP_1743), *actP*-II (PP_2797), and *actP*-III (PP_3272) (Figure 3a). To determine their role in acetate uptake, single-, double-, and triple-deletion mutants lacking the respective genes (seven in total) were created, and their phenotype was investigated during growth on acetate. None of the mutants showed a phenotype, with all growing similarly to the wild type on acetate (Figure 5b). Testing growth of *P. putida* KT2440 Δ*actP*-I Δ*actP*-II Δ*actP*-III across different acetate concentrations (0.6 -10 g L^-1^) revealed neither the mutant nor the wild type were able to grow on concentrations higher than 6 g L^-1^ acetate. Interestingly, the deletion mutant showed shorter lag phases than the wild type on all tested concentrations (Figure 3c), suggesting that the absence of these putative acetate transporters provided an advantage to the cell rather than a drawback. This indicates that acetate likely does not enter the cell through these permeases at concentrations between 0.6 and 6 g/L. However, it remains possible that at lower concentrations, these transporters play a role, as seen in *Geobacter*, where transcript levels of acetate permease-like genes increased as acetate concentration decreased (Elifantz et al. 2010).

**Figure 3:**
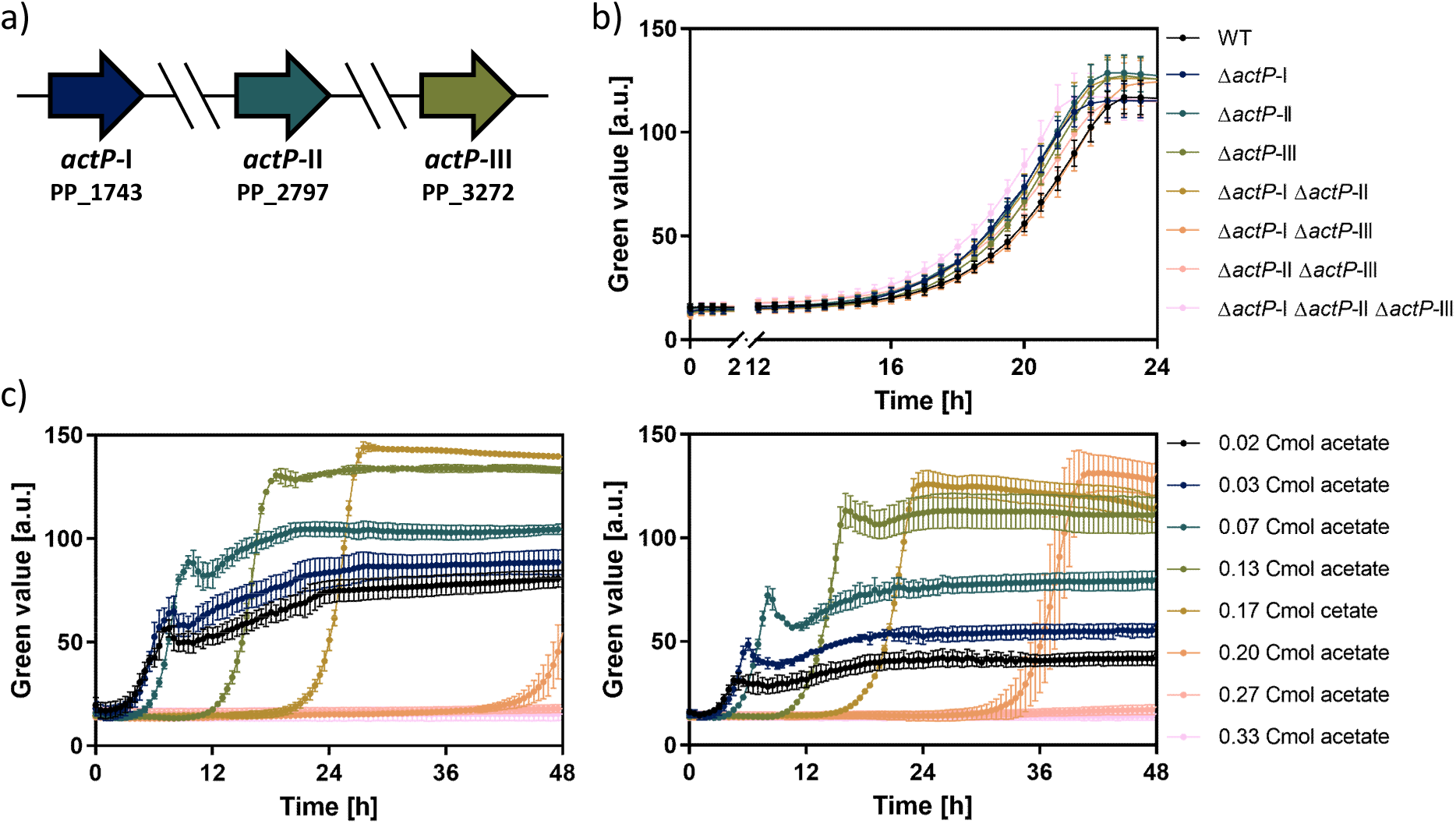
Acetate permeases in *P. putida* KT2440. a) Genomic orientation of three *actP*-genes in *P. putida* KT2440. b) Growth of *P. putida* KT2440 mutants deficient in *actP*-genes on 5 g L^-1^ acetate. c) Cultivation of *P. putida* KT2440 (left) and *P. putida* KT2440 Δ*actP*-I Δ*actP*-II Δ*actP*-III (right) on different acetate concentrations. Error bars represent the standard deviation of three biological replicates.

To determine if *P. putida* KT2440 contains other homologs of acetate permeases that might compensate for the function in the Δ*actP* triple-KO mutant, a BLASTp search (Altschul et al. 1990) was performed to identify proteins with a similar amino acid sequence. Two further genes showing 23.8 and 25.8 % identity to the amino acid sequence of ActP-III were identified: PP_4946, a sodium/proline symporter, and PP_4524, a sodium/solute symporter. Deletion mutants of both genes, individually and in combination with the *actP*-deletions, were tested for growth on acetate. All mutants exhibited growth rates comparable to the wild type on 5 g L^-1^ acetate (Figure S 1). All of the tested mutants were still able to produce biomass from low (0.5 g L^-1^) and higher (5 g L^-1^) acetate concentrations, suggesting none of the deleted genes was essential for acetate uptake.

These findings align with previous studies showing that the *actP* genes are not upregulated upon growth on acetate as sole carbon source (Henriquez and Jung 2021). RNA-seq data collected in this study (see section 3.9) did not reveal a significantly higher expression of any investigated genes coding for putative acetate transporters during growth on acetate compared to growth on glucose, confirming previous data.

To explore whether ActP transporters facilitate uptake of other organic acids, the *P. putida* KT2440 mutants *P. putida* KT2440 Δ*actP* I-III and *P. putida* KT2440 Δ*actP* I-III ΔPP_4524 ΔPP_4946 were grown on propionate (C3), hexanoate (C6), octanoate (C8), decanoate (C10), succinate (C4), malate (C4), and lactate (C3), as well as glucose and acetate (C2) as a control. The two mutants were still able to grow on all the tested substrates without exhibiting a major growth deficit (Figure S 2). Accordingly, either none of the tested organic acids was the target molecule of the deleted transporters, or there is significant promiscuity among other transporters that could fulfill this role.

### 3.5 Identification of potential bottlenecks in the acetate metabolism

It was investigated whether the second step in acetate metabolism, the activation of intracellular acetate to acetyl-CoA constitutes a bottleneck in acetate metabolization.

Literature describes two ways of acetate activation: the one-step acetyl-CoA synthetase (ACS) and the two-step AckA-PTA, which both rely on ATP. While *E. coli* features both, *P. putida* KT2440 lacks the *ackA* gene, relying solely on ACS (Figure 1a). In *P. putida* KT2440, two genes (PP_4487, *acsA*-I, and PP_4702, *acsA*-II) are predicted to encode acetyl-CoA synthetases. Furthermore, at least three genes are predicted to code for acyl-CoA synthetases acting on organic acids of different lengths: PP_3458 (*acs*), PP_2213, and PP_3724. Single deletion mutants were constructed to identify the gene required for acetate growth in *P. putida* KT2440.

While four of the mutants exhibited the same growth behavior as the wild type, ΔPP_4487 exhibited a prolonged lag phase and a reduced growth rate (0.20 h^-1^ compared to 0.31-0.36 h^-1^), indicating that *acsA*-I is required for growth on acetate (Figure 4a). After more than 24 hours when the mutant starts growing, the strain likely re-routes the metabolism and uses a different homolog for the activation of acetate. In order to test if activation of acetate via PP_4487 was a bottleneck, the gene was overexpressed. A second copy of PP_4487 was integrated into PP_0340, a suitable landing pad for heterologous gene expression in *P. putida* KT2440 (Köbbing et al. 2024). Growth of the overexpression mutant was compared to growth of the wild type on acetate, as well as on glucose. While the overexpression hardly affected the growth rate on glucose (0.29 h^-1^ for the wild type compared to 0.32 h^-1^ for the mutant), it did show positive effects on acetate (Figure 4b). The lag phase was reduced by approximately six hours and the growth rate was increased from 0.25 h^-1^ to 0.31 h^-1^, strengthening the hypothesis that PP_4487 is involved in acetate metabolization under the chosen growth conditions and that the activation of acetate constitutes a bottleneck.

**Figure 4:**
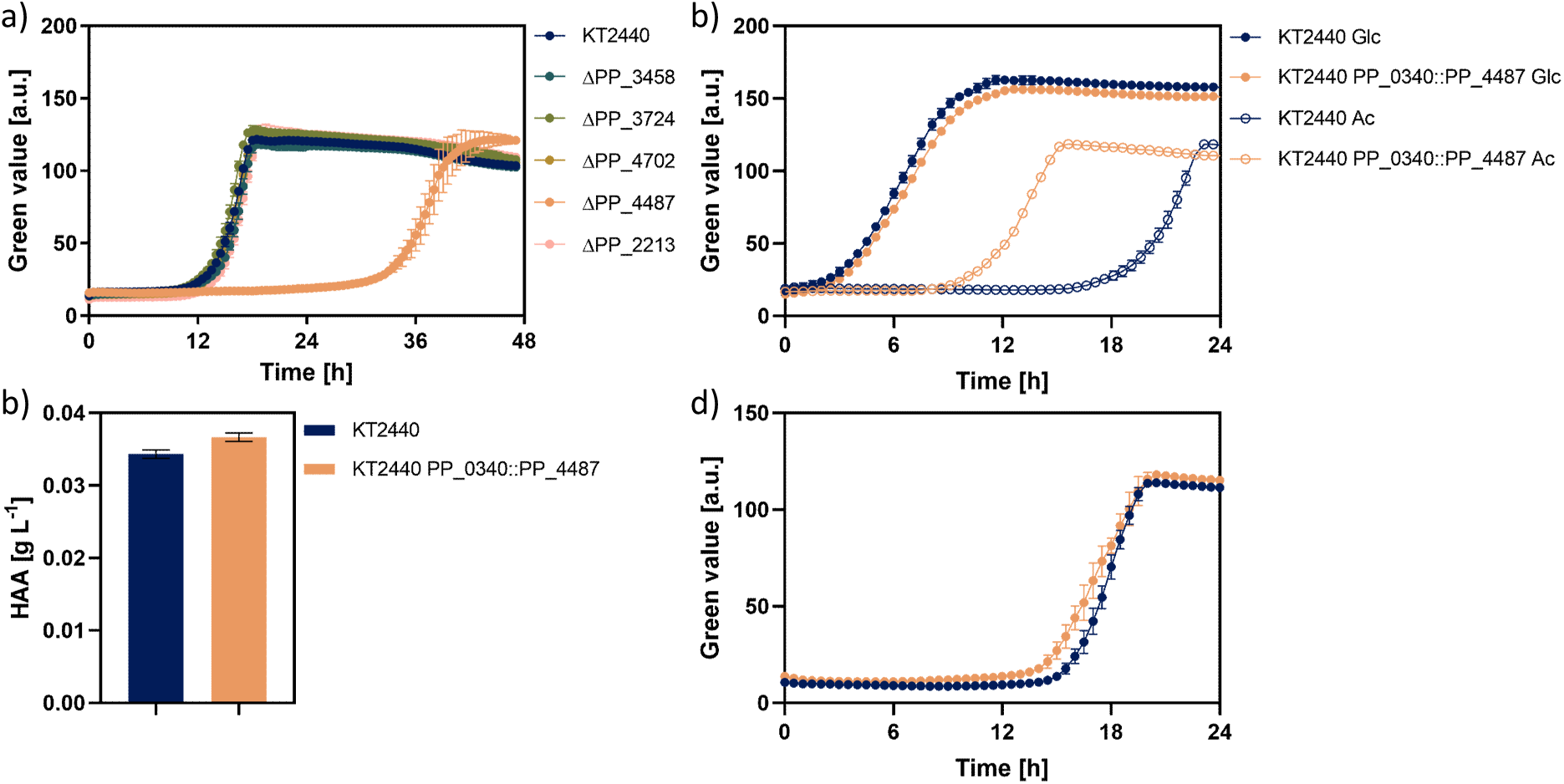
Investigating the acetate catabolic network in *P. putida* KT2440. a) Growth Profiler cultivation of single deletion mutants deficient in one of five ac(et)yl-CoA synthetase encoding genes of *P. putida* KT2440 on MSM 5 g L^-1^ acetate and b) *P. putida* KT2440 and *P. putida* KT2440 PP_0340::PP_4487 on MSM + glucose or MSM + acetate. c) Final HAA titers of *P. putida* KT2440 HAA and *P. putida* KT2440 PP_0340::PP_4487 HAA on MSM + acetate. d) Growth Profiler cultivation of *P. putida* KT2440 (blue) and *P. putida* KT2440 *att*Tn7::P_ffg_-*aceA*-*glcB* (orange). Error bars represent standard deviation of three biological replicates.

**Figure 5:**
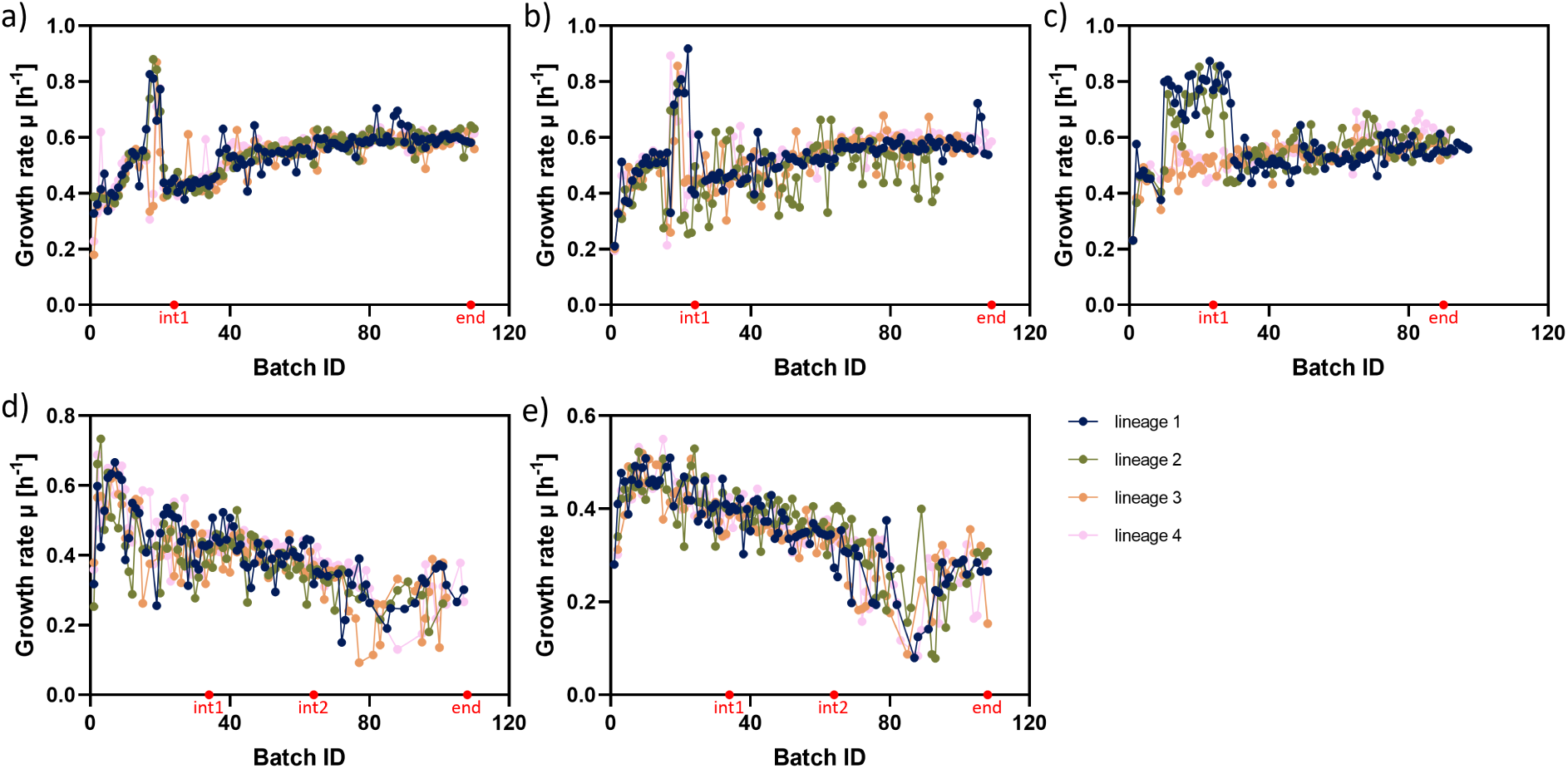
Growth rates during laboratory evolution of *P. putida* strains. a) *P. putida* KT2440 ALE, b) *P. putida* KT2440 GR18 ALE, c) *P. putida* KT2440 GR18a ALE, d) *P. putida* KT2440 TALE, b) *P. putida* KT2440 GR18 TALE. Colors (blue, green, orange and pink) indicate independent lineages. Glycerol stocks were made as indicated by red dots. int1 = intermediate 1, int2 = intermediate 2, end = endpoint. High growth rates at batch #17-20 are artifacts of an incorrectly prepared medium with a reduced acetate concentration.

Since the activation of acetate yields acetyl-CoA, overexpressing this step should result in faster acetyl-CoA generation. To assess if this surplus of available acetyl-CoA could enhance formation of an acetyl-CoA-derived product, HAA biosynthesis was investigated in the *acsA*-I overexpression mutant *P. putida* KT2440 PP_0340::PP_4487. The mutant showed a growth advantage, mainly visible by a reduced lag phase (data not shown), and a slightly higher HAA titer than the wild type (Figure 4c), suggesting that overexpression of *acsA*-I can enhance both growth and acetyl-CoA-derived product formation.

The glyoxylate shunt is essential for anaplerotic and gluconeogenic reactions during growth on C2 substrates like acetate, as mutants lacking this pathway cannot grow on such substrates. Furthermore, genome-scale metabolic modelling (*i.e.*, Flux Balance Analysis, FBA) carried out by Ziegler et al. (2023) predicted significantly higher fluxes through the TCA cycle and glyoxylate shunt during growth on acetate versus glucose, with a 6.7-fold increase in flux through the isocitrate lyase. In addition, RNA-seq data revealed higher mRNA levels for the genes coding for isocitrate lyase and malate synthase G (discussed in section 3.9), indicating that the demand on the two reactions catalyzed by enzymes encoded by *aceA* and *glcB* is increased and might even constitute a bottleneck on the C2 compound. To test this, a second copy of *aceA* and *glcB* was integrated into the *att*Tn7-site *P. putida* KT2440 under the control of the stacked synthetic promoter ffg (Köbbing et al. 2020). However, overexpression showed no improvement in growth on acetate (Figure 4d), indicating these reactions are not limiting steps in the wild-type background.

### 3.6 Adaptive laboratory evolution improves growth performance on acetate

An effective non-rational approach, which is orthogonal to the rational engineering approach presented above, was chosen as the next step to improve the growth performance of *P. putida* KT2440 on acetate: adaptive laboratory evolution (ALE). Due to constant selective pressure, the occurrence of beneficial mutations resulting in superior growth on acetate was expected.

ALE experiments of *P. putida* KT2440 were performed in MSM with 5 g L^-1^ acetate in four parallel independent replicate evolution experiments. Additionally, two genome-reduced (GR) strains, *i.e*., *P. putida* KT2440 GR18 and *P. putida* KT2440 GR18a, were used for the ALE experiment, both with four independent replicates as well, to compare mutational behavior to the wild-type strain. The two GR strains lack large genomic regions coding for flagellar machinery, biofilm-forming components (the exopolysaccharides alginate, cellulose, Pea, and Peb, and the extracellular adhesins LapA and LapF), the formation of the side product pyoverdines, and the internal storage PHA to streamline the metabolism and free up resources for the production of a product of choice. Since many energy- and resource-demanding reactions were deleted, the question arose if this would influence the mutational behavior of *P. putida* KT2440.

While wild type *P. putida* KT2440 exhibited an initial growth rate of 0.43 h^-1^, *P. putida* KT2440 GR18 exhibited an initial growth rate of 0.45 h^-1^. After ALE, the growth rate of adapted *P. putida* KT2440 strains increased to 0.58 - 0.62 h^-1^ after 109 - 114 transfers. The genome reduced strain GR18 reached a growth rate of 0.57 h^-1^ after 105 transfers, while *P. putida* KT2440 GR18a, which exhibited an initial growth rate of 0.46 h^-1^, achieved this growth rate after 93 transfers (Table 2). Thus, all three starting strains used reached comparable growth rates on acetate after undergoing ALE, independent of the initial genotype.

Besides improving the growth rate during ALE, acetate tolerance was improved in a separate tolerance ALE (TALE) experimental protocol (Mohamed et al. 2017). Unlike ALE, where the acetate concentration remained constant, the substrate concentration was increased throughout the TALE experiments. When the growth rate recovered on an increasing concentration, the concentration was increased again, forcing an adaptation to even higher acetate concentrations. *P. putida* KT2440 and *P. putida* KT2440 GR18, initially grown on 5 g L^-1^ acetate, tolerated 16 g L^-1^ after 103 - 116 transfers (Table 3).

**Table 3:** Growth rates of each individually evolved lineage of *P. putida* KT2440 (ALE ID 13-16) and *P. putida* KT2440 GR18 (ALE ID 17-20) TALE experiment, including the number of flasks, generations, and acetate concentration. *Average of the initial growth rate for all four lineages. **Average of the five last flasks.

| ALE ID | Number of transfers | Number of generations | Initial growth rate [h<sup>-1</sup>]* | Final growth rate [h<sup>-1</sup>]** | Start concentration [g L<sup>-1</sup>] | Final acetate concentration [g L<sup>-1</sup>] |
| --- | --- | --- | --- | --- | --- | --- |
| 13 | 108 | 534 | 0.43 ± 0.02 | 0.28 ± 0.02 | 5 | 16 |
| 14 | 103 | 512 |  | 0.26 ± 0.01 |  | 16 |
| 15 | 103 | 509 |  | 0.36 ± 0.01 |  | 16 |
| 16 | 11 | 552 |  | 0.35 ± 0.01 |  | 16 |
| 17 | 109 | 541 | 0.45 ± 0.02 | 0.27 ± 0.01 |  | 16 |
| 18 | 116 | 573 |  | 0.30 ± 0.02 |  | 16 |
| 19 | 108 | 531 |  | 0.31 ± 0.01 |  | 16 |
| 20 | 110 | 542 |  | 0.28 ± 0.01 |  | 16 |

The growth rate of the evolved population was monitored throughout the experiment (Figure 6). In the ALE experiments, *P. putida* KT2440, *P. putida* KT2440 GR18, and *P. putida* KT2440 GR18a exhibited increasing growth rates until stabilization (Figure 6b, c). In TALE, however, growth rates of *P. putida* KT2440 and *P. putida* KT2440 GR18 decreased with increasing acetate concentration (Figure 6d, e). While adaptation to acetate during the ALE experiment resulted in improved growth performance, probably due to rechanneling of the metabolism, TALE enabled adaptation to a higher concentration at the cost of the growth rate, probably due to improved tolerance mechanisms, highlighting the distinct selection pressures of the two approaches of evolutionary engineering.

**Figure 6:**
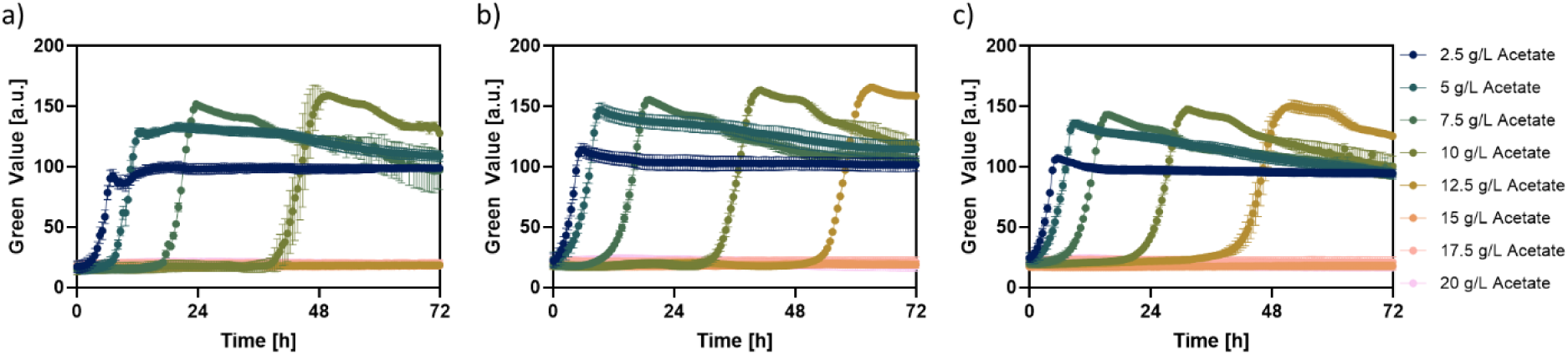
Growth of evolved strains on different acetate concentrations. a) *P. putida* KT2440, b) *P. putida* KT2440 ALE and c) *P. putida* KT2440 TALE. Cultivation was performed in 24-well microtiter plates with transparent bottom with 1.5 ml filling volume in a growth profiler. Green value was determined as a measure of culture turbidity, *i.e*., biomass formation. Error bars represent standard deviation of three biological replicates.

After the ALE experiments, the resulting populations were cryopreserved, and single colonies were screened for superior growth on acetate. Among the different lineages and isolates, the best performer of *P. putida* KT2440 ALE, *i.e.*, ALE ID 4, flask 114, isolate 6, R1, and *P. putida* KT2440 TALE, *i.e*., ALE ID 13, flask 108, isolate 7, R1, was selected during a screening in minimal medium with acetate as sole carbon source performed in a Growth Profiler (Enzyscreen, Heemstede, The Netherlands). To assess the evolved mutants of *P. putida* KT2440, the best ALE isolate (improved growth rate) and the best TALE isolate (improved tolerance) were characterized on different acetate concentrations compared to the initial strain.

The initial strain was impacted by relatively low acetate concentrations and failed to grow on concentrations exceeding 10 g L^-1^ within 72 hours, while both evolved strains were able to grow on acetate concentrations up to 12.5 g L^-1^ within 72 h (Figure 7). Even at 2.5 g L^-1^, the evolved strains showed a clear growth advantage. Lag phases and growth rates were determined as a measure of tolerance towards acetate. While the lag phase is an indicator of the resistance against acetate toxicity, the growth rate represents the catabolic activity (Mohamed et al. 2020).

**Figure 7:**
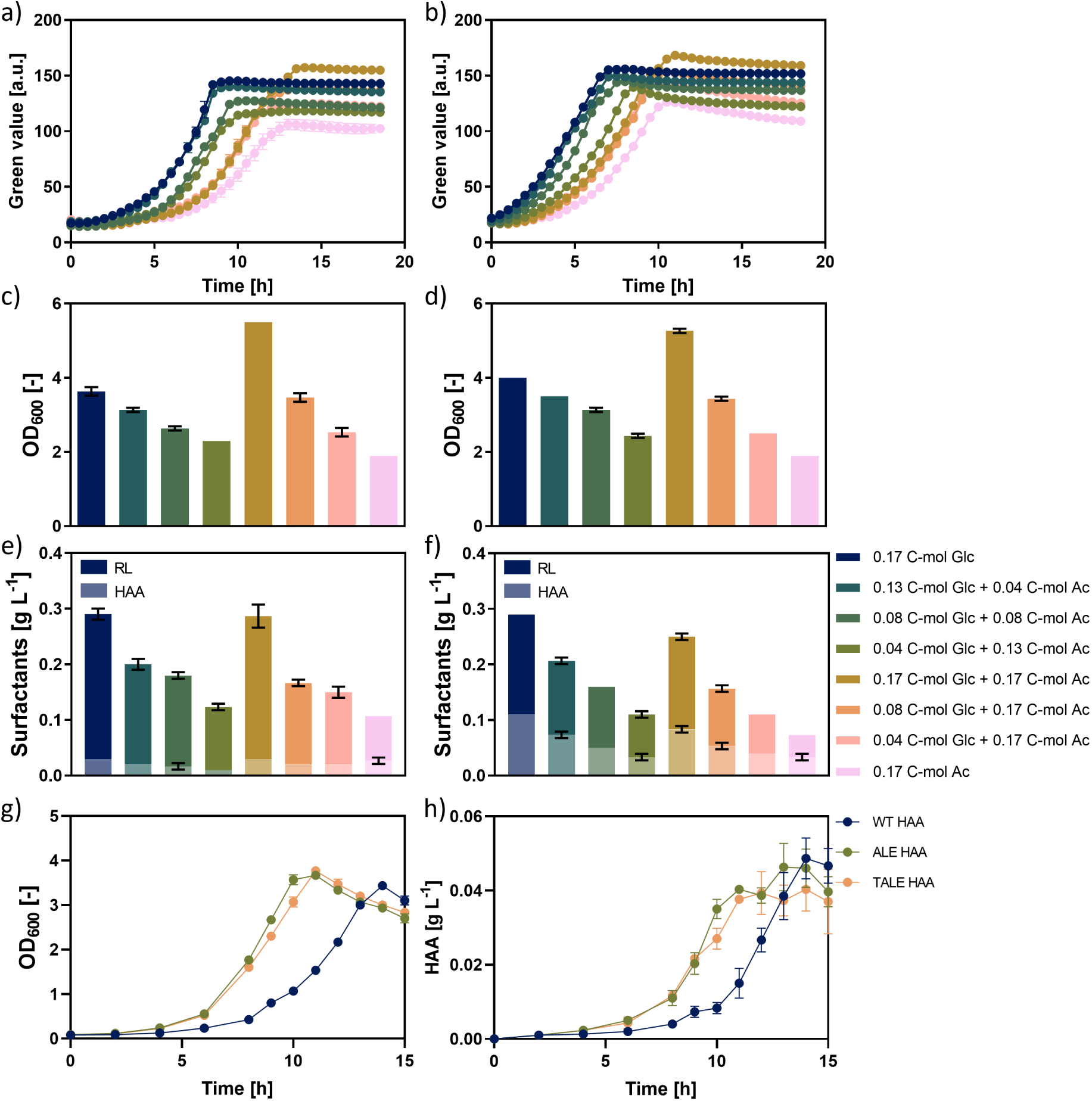
Rhamnolipid and HAA production with evolved strains of *P. putida* KT2440. a)-f) show results of a Growth Profiler cultivation of SK4 (left) and *P. putida* KT2440 ALE RL (right) on different ratios of glucose and acetate. a) + b) Growth, c) + d) final OD and e) + f) final surfactant titer obtained during cultivation. g) + h) show shake flask cultivation of *P. putida* KT2440 *att*Tn7::P_ffg_-*rhlA* (blue), *P. putida* KT2440 ALE *att*Tn7::P_ffg_-*rhlA* (green) and *P. putida* KT2440 TALE *att*Tn7::P_ffg_-*rhlA* (orange) on 5 g L^-1^ acetate. g) Optical density as a measure of biomass formation and h) HAA concentration. Error bars represent standard deviation of three biological replicates.

*P. putida* KT2440 TALE (ALE ID 13, flask 108, isolate 7) depleted acetate first and exhibited the shortest lag phases (Table 4). However, *P. putida* KT2440 ALE (ALE ID 4, flask 114, isolate 6) exhibited higher growth rates, as expected given their distinct evolutionary pressures. Overall, the growth rates were found to decrease with increasing acetate concentration.

**Table 4:** Growth rates and lag phases exhibited by *P. putida* KT2440 WT, *P. putida* KT2440 ALE, and *P. putida* KT2440 TALE on different acetate concentrations from 2.5 g L^-1^ to 20 g L^-1^. n.d.: No growth detected within 72 h.

| Acetate concentration<br>[g L<sup>-1</sup>] | WT |  | ALE |  | TALE |  |
| --- | --- | --- | --- | --- | --- | --- |
| | Growth rate<br>$\mu$ [h<sup>-1</sup>] | Lag phase<br>[h] | Growth rate<br>$\mu$ [h<sup>-1</sup>] | Lag phase<br>[h] | Growth rate<br>$\mu$ [h<sup>-1</sup>] | Lag phase<br>[h] |
| 2.5 | 0.36 | 1.5 | 0.34 | 0.0 | 0.31 | 0.0 |
| 5.0 | 0.43 | 4.0 | 0.25 | 1.5 | 0.26 | 1.5 |
| 7.5 | 0.36 | 13.1 | 0.22 | 7.1 | 0.24 | 2.5 |
| 10.0 | 0.28 | 35.0 | 0.28 | 28.2 | 0.23 | 14.6 |
| 12.5 | n.d. | n.d. | 0.26 | 46.7 | 0.21 | 30.6 |
| 15.0 | n.d. | n.d. | n.d. | n.d. | n.d. | n.d. |
| 17.5 | n.d. | n.d. | n.d. | n.d. | n.d. | n.d. |
| 20.0 | n.d. | n.d. | n.d. | n.d. | n.d. | n.d. |

### 3.7 Acetate-to-glucose ratio in mixed substrate impacts share of HAAs in surfactant mixture produced by evolved strains

After confirming the superior growth of the evolved strains, the potential to translate this improved performance into producing an acetyl-CoA-derived secondary metabolite was investigated. The chosen product was the biosurfactant rhamnolipid. Rhamnolipid biosynthesis genes *rhlA* and *rhlB* were integrated into the ALE strain to compare biosurfactant production to the wild type. Biosurfactant production was determined on different glucose-to-acetate ratios, as well as on glucose and acetate as sole carbon sources, whereas the total carbon amount or the acetate amount was constant. While biomass production of the evolved strain was superior to that of the wild type, just as observed for the non-producing strains (Figure 8a-d), biosurfactant production of the evolved strain was slightly reduced compared to the wild type under all conditions, except on glucose alone, where titers were comparable (Figure 8, f). Interestingly, the evolved strain produced a higher proportion of HAAs, reaching almost 50 % on acetate as the sole carbon source. This phenomenon, which was much more pronounced in the evolved strain, is likely caused by the altered metabolism of the evolved strains, hinting to a reduced gluconeogenic flux.

**Figure 8:**
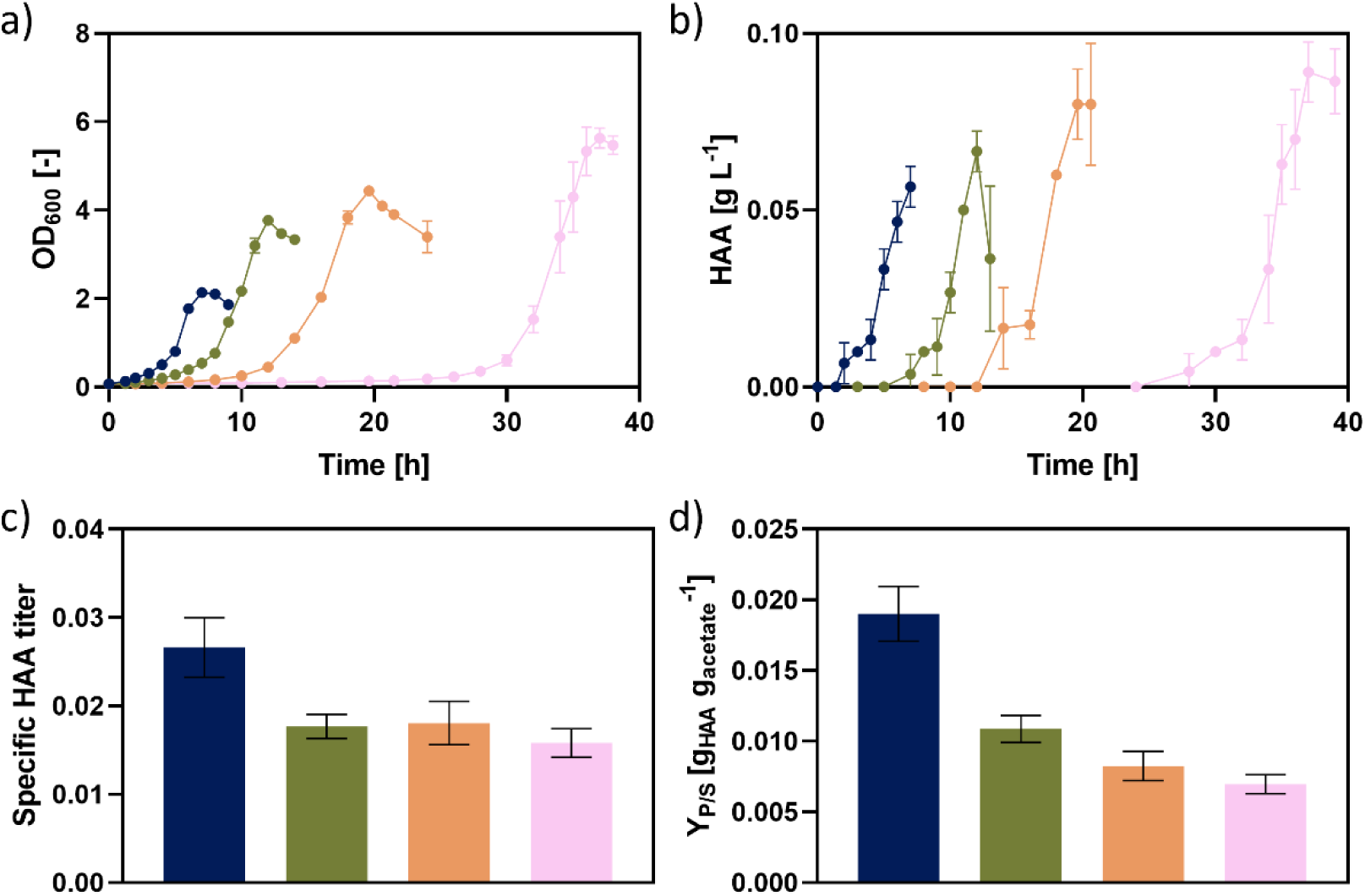
Cultivation of *P. putida* KT2440 TALE *att*Tn7::P_ffg_-*rhlA* on 2.5 g L^-1^ (blue), 5 g L^-1^ (green), 7.5 g L^-1^ (orange), and 10 g L^-1^ acetate (pink). a) Biomass formation, b) HAA production, c) specific HAA titer and d) product yield. Cultivation was performed in 50 ml shake flasks with 10 % filling volume at 30°C and 250 rpm. Error bars represent standard deviation of three biological replicates.

Since the evolved strain reached higher HAA concentrations in the product mixture, HAA producers of the ALE and TALE strains were created by integrating only the first gene of the rhamnolipid biosynthesis pathway, the acyltransferase *rhlA*. HAA production was compared to the wild type on 5 g L^-1^ acetate. Although final biomass levels were similar across all strains, HAA titers reached by the ALE and TALE strains were slightly lower than the titer reached by the wild type, reaching 94 % and 82 %, respectively (Figure 8g, h). Congener distribution was consistent across strains. Although the HAA concentrations reached in this experiment were generally low, it is not surprising that the evolved strains exhibit a lower product formation than the wild type since the strains were not evolved towards higher product formation. Overall, the evolved strains show a clear advantage regarding biomass formation and lag phase while still reaching comparable product titers, at least the ALE strain. All in all, the evolved strains reach the same space-time yields as the wild type (3.5 mg L^-1^ h^-1^ for ALE and 3 mg L^-1^ h^-1^ for TALE compared to 3.5 mg L^-1^ h^-1^ for WT).

To investigate HAA production of *P. putida* KT2440 TALE HAA at higher acetate concentrations, the strain was cultivated on 2.5 g L^-1^, 5 g L^-1^, 7.5 g L^-1^, and 10 g L^-1^ acetate. As seen in previous experiments for the non-producing strain, the lag phase increased, and the growth rate decreased with increasing acetate concentrations (Figure 9a). Both, final biomass concentration, and HAA titers, increased with higher acetate concentrations (Figure 9b), although the increase in HAA titers did not match the increase in carbon availability. The product yield declined with increasing acetate concentration from 19 mg_HAA_ g_substrate_^-1^ on 2.5 g L^-1^ acetate to 10 mg_HAA_ g_substrate_^-1^ on 5 g L^-1^, 8 mg_HAA_ g_substrate_^-1^ on 7.5 g L^-1^ and 7 mg_HAA_ g_substrate_^-1^ on 10 g L^-1^ acetate. The data show that increasing the initial acetate concentration in the medium is not an optimal way to increase product titers. Since the consumption of acetate results in a rise in the pH of the cultivation broth, one issue contributing to the reduced product yields might be the higher final pH from higher acetate starting concentrations, which influences cell performance. Although the strain was evolved towards higher acetate tolerance, the results point out the limits of shake flask cultivations and highlight the need for a bioprocess with a suitable feeding strategy to avoid pH-related issues.

**Figure 9:**
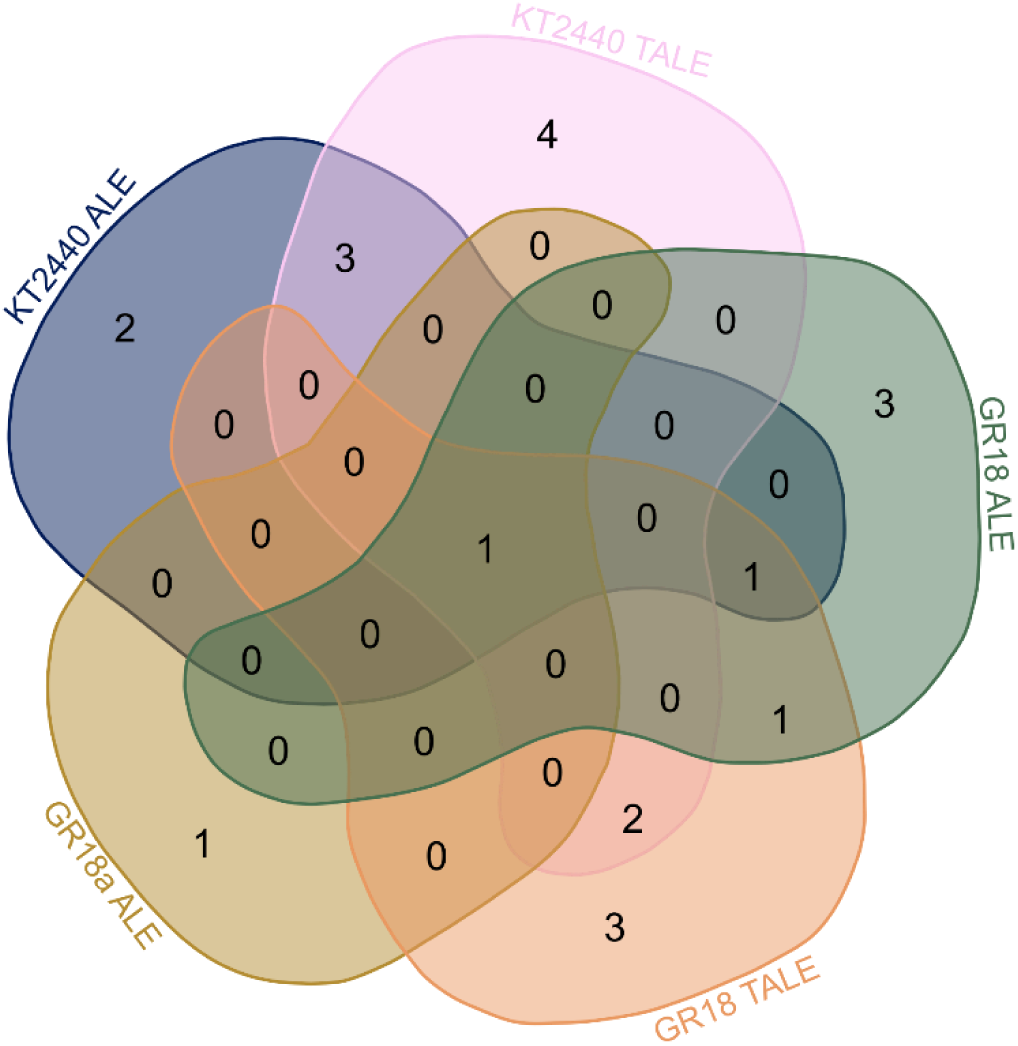
Venn diagram showing converged mutations in ALE and TALE experiments of *P. putida* KT2440, *P. putida* KT2440 GR18, and *P. putida* KT2440 GR18a. Affected genes were identified using whole genome sequencing. Venn diagram was calculated using https://bioinformatics.psb.ugent.be/webtools/Venn/.

### 3.8 Whole genome sequencing revealed mutations underlying improved growth on acetate

Whole genome sequencing was conducted to pinpoint mutations that enabled the improved phenotypes of the evolved strains. Therefore, single colonies with superior growth on acetate were isolated and their genomic DNA was sequenced. A total of 76 isolates were sequenced: for *P. putida* KT2440 ALE, *P. putida* KT2440 GR18 ALE, and *P. putida* KT2440 GR18a ALE two isolates of the intermediate 1 (population after 24 transfers) and one isolate of the endpoint of each of the four replicates were sequenced. For *P. putida* KT2440 TALE and *P. putida* KT2440 GR18 TALE two isolates of intermediate 1 (population after 34 transfers) and intermediate 2 (population after 64 transfers) each, and one isolate of the endpoint of all four replicates were sequenced. Sampling points during the course of the cultivation are depicted in Figure 6. Starting strains were sequenced in order to filter out starting strain-specific mutations. A complete list of mutations can be found in Table S 2. Since these sequencings revealed hundreds of mutations, only converged mutations, *i.e*., mutations occurring in more than two independent ALE or TALE experiments, were considered for further analysis (Table 5). Six of the 23 converged mutations were found in intergenic, non-coding regions.

**Table 5:** Converged mutations found in (T)ALE experiments of *P. putida* strains on acetate. *Identified according to Winsor et al. (2016). †Selected as targets for reverse engineering. SNP = single nucleotide polymorphism, DEL = deletion, INS = insertion.

| Gene name<br>(Locus ID) | Gene Product* | Experiment | Mutation type<br>(unique counts) |
| --- | --- | --- | --- |
| *rpoD* (PP_0387) | RNA polymerase sigma 70 factor | KT2440 ALE | SNP (2) |
| *gacS* (PP_1650)† | sensor protein GacS | KT2440 ALE, KT2440 TALE, GR18 ALE, GR18 TALE, GR18a ALE | SNP (9), DEL (8), INS (3) |
| *uvrY* (PP_4099)† | BarA/UvrY two-component system response regulator | KT2440 ALE, GR18 ALE, GR18 TALE | SNP (5), DEL (1) |
| *sdhC* (PP_4193),<br>*gltA* (PP_4194)<br>(intergenic) | succinate dehydrogenase membrane b-556 subunit/ citrate synthase | KT2440 ALE | SNP (1) |
| *fleQ* (PP_4373)† | transcriptional regulator FleQ | KT2440 ALE, KT2440 TALE | SNP (6), DEL (3), INS (1) |
| *recB* (PP_4673) | Chi activated ATP-dependent DNA helicase/dsDNA/ssDNA exonuclease | KT2440 ALE, KT2440 TALE | SNP (2) |
| PP_4743 | hypothetical protein | KT2440 ALE, KT2440 TALE | DEL (2) |
| *oprQ* (PP_0268) | outer-membrane porin D | KT2440 TALE | SNP (2), DEL (1) |
| *livM* (PP_1139) | branched-chain amino acid ABC transporter permease | KT2440 TALE | DEL (2) |
| *livH* (PP_1140) | branched-chain amino acid ABC transporter permease | KT2440 TALE | DEL (1) |
| *cbrB* (PP_4696),<br>*pcnB* (PP_4697)<br>(intergenic) | response regulator, CbrB/ poly(A) polymerase | KT2440 TALE | DEL (1) |
| *crc* (PP_5292)† | catabolite repression control protein | KT2440 TALE, GR18 TALE | SNP (3), DEL (2) |
| *aruC* (PP_0372),<br>*yqaE* (PP_0373)<br>(intergenic) | acetylornithine aminotransferase/ membrane protein | GR18 ALE | SNP (1) |
| PP_1878 | hypothetical protein | GR18 ALE, GR18 TALE | SNP (2) |
| *gllP* (PP_2517) | porin-like protein | GR18 ALE | SNP (1) |
| PP_3486,<br>PP_3487<br>(intergenic) | cytochrome c/ hypothetical protein | GR18 ALE | SNP (1) |
| *rpoC* (PP_0448) | DNA-directed RNA polymerase subunit beta'Add | GR18 TALE | SNP (2) |
| *pfeS*-II<br>(PP_1652)† | histidine kinase | GR18 TALE | SNP (5) |
| PP_RS09190,<br>PP_RS09195,<br>PP_RS09200,<br>PP_RS09205,<br>PP_RS09210,<br>PP_RS09215,<br>PP_RS09220,<br>PP_RS09225,<br>PP_RS09230 | hypothetical protein, hypothetical protein, HAD family hydrolase, acylneuraminate cytidyltransferase family protein, aldolase catalytic domain-containing protein, glycosyltransferase family 2 protein, glycosyltransferase family 2 protein, DUF4214 domain-containing protein, hypothetical protein | GR18 TALE | DEL (1) |
| *lon*-II (PP_2302),<br>*hupB* (PP_2303)<br>(intergenic) | DNA-binding ATP-dependent protease/ DNA binding regulator subunit beta B | GR18 TALE | DEL (1) |
| *hupB* (PP_2303) | DNA binding regulator subunit beta B | GR18 TALE | SNP (3) |
| *uvrY*, PP_4100<br>(intergenic) | BarA/UvrY two-component system response regulator/Cro/CI family transcriptional regulator | GR18 TALE | SNP (1) |

Most of the affected genes were unique to individual experiments. However, three affected genes were shared between *P. putida* KT2440 ALE and *P. putida* KT2440 TALE. One affected gene was common to both *P. putida* KT2440 TALE and *P. putida* KT2440 GR18 TALE, and another affected gene was found in sequencing results of *P. putida* KT2440 GR18 ALE and *P. putida* KT2440 GR18 TALE. Additionally, one gene was affected in *P. putida* KT2440 ALE, *P. putida* KT2440 GR18 ALE, and *P. putida* KT2440 GR18 TALE. Only one gene was found to be affected in all five experiments (Figure 10).

**Figure 10:**
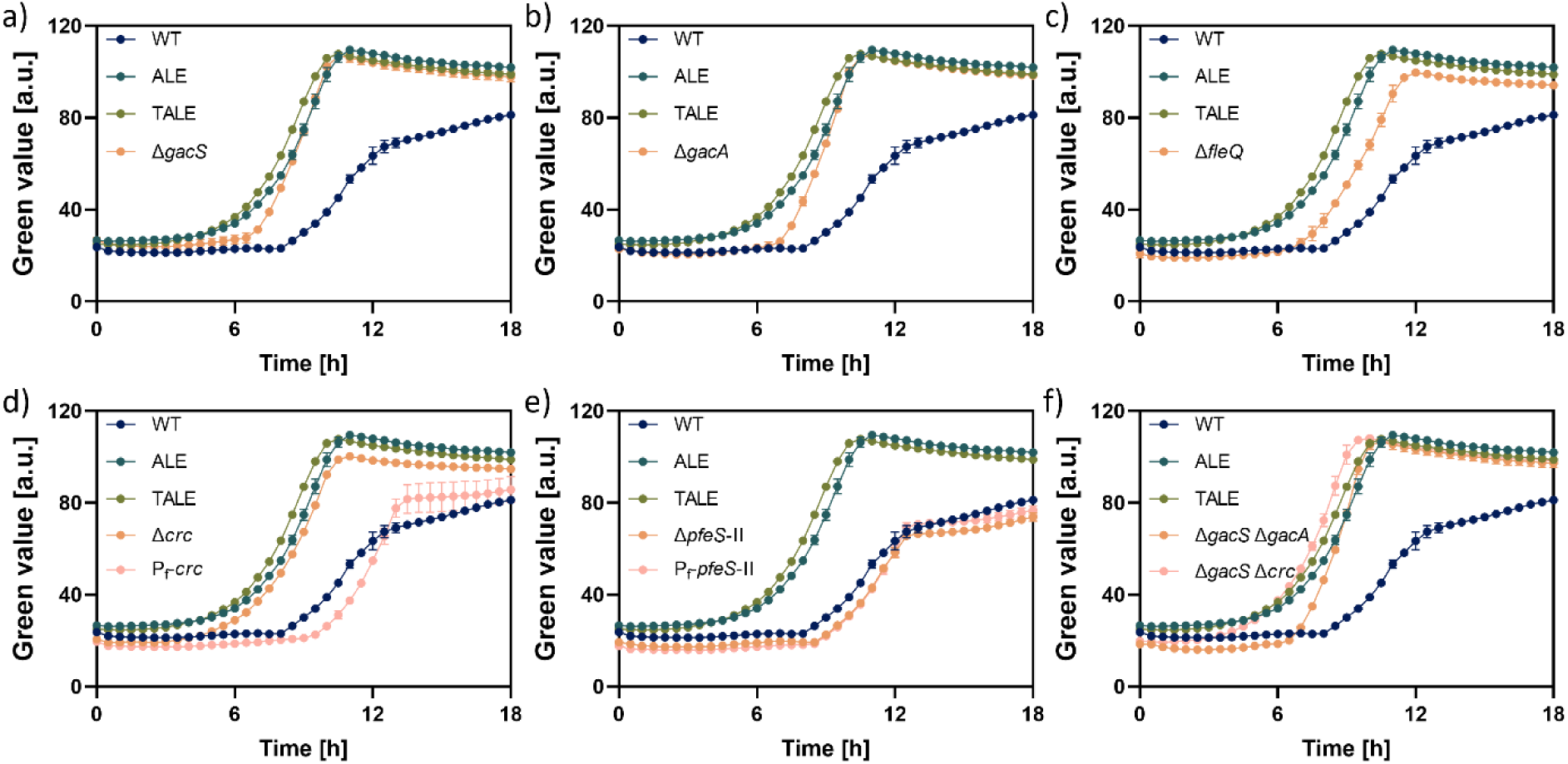
Growth of *P. putida* KT2440, *P. putida* KT2440 ALE, *P. putida* KT2440 TALE, and a) *P. putida* KT2440 Δ*gacS*, b) *P. putida* KT2440 Δ*gacA*, c) *P. putida* KT2440 Δ*fleQ*, d) *P. putida* KT2440 Δ*crc*, and *P. putida* KT2440 *att*Tn7::P_f_-*crc*, e) *P. putida* KT2440 Δ*pfeS-*II, and *P. putida* KT2440 *att*Tn7::P_f_-*pfeS*-II, and f) *P. putida* KT2440 Δ*gacS* Δ*gacA* and *P. putida* KT2440 Δ*gacS* Δ*crc* on 5 g L^-1^ acetate. Cultivation was performed in 24-well microtiter plates with transparent bottom with 1.5 ml filling volume in a growth profiler. Green value was determined as a measure of culture turbidity, *i.e*., biomass formation. Error bars represent standard deviation of three biological replicates.

Analysis of the converged mutations identified five key genes repeatedly affected across experiments, making them promising targets for reverse engineering: PP_1650 (*gacS*), PP_4099 (*uvrY*), PP_4373 (*fleQ*), PP_5292 (*crc*), as well as PP_1652 (*pfeS*-II). Mutations in PP_5292 were found in both TALE experiments and emerged only in the late phase of the experiments, *i.e*., at high acetate concentrations, suggesting a role in tolerance towards acetate. Similarly, mutations in PP_1652 exclusively appeared in later stages of the GR18 TALE experiment and, thus, also indicating an involvement in tolerance towards acetate.

PP_1650 (*gacS*) encodes GacS, the sensor protein in the GacS/GacA two-component system involved in the regulation of several physiological processes, including the synthesis of secondary metabolites and in biofilm formation (Workentine et al. 2009), as well as in flagellar formation (Kim et al. 2014). Sequencing revealed the presence of several nonsense mutations and deletions, likely disrupting the function of GacS. GacS was also found as a target of mutation during ALE experiments of *P. putida* KT2440 on other compounds, including xylose (Lim et al. 2021), *p*-coumaric and ferulic acid (Mohamed et al. 2020), and ionic liquids (Lim et al. 2020). The second part of the two-component system, GacA/UvrY, encoded by PP_4099, was also affected in the ALE experiments, as seen in prior studies on *p*-coumaric and ferulic acid (Mohamed et al. 2020), as well as ionic liquids (Lim et al. 2020). PP_4373 encodes the transcriptional regulator FleQ, which regulates genes involved in flagella and biofilm formation (Blanco-Romero et al. 2018). Mutations in *fleQ* included nonsense, and frameshift mutations as well as deletions of several base pairs, likely abolishing the function of this gene. Mutations disrupting *fleQ* were also previously observed in ALE experiments with *P. putida* KT2440 on ethanol (Bator et al. 2020), on syringate (Mueller et al. 2022), as well as on ferulic acid and glucose (Mohamed et al. 2020). PP_5292 encodes a catabolite repression control (CRC) protein. From the sequencing data, it cannot be stated whether the function of this gene is disrupted or upregulated by the mutations. PP_1652 (*pfeS*-II) encodes a sensor histidine kinase. Since *gacS*, *gacA,* and *fleQ* apparently were disrupted during ALE experiments, deletion mutants of the respective genes were generated and cultivated on acetate. All three deletion mutants exhibited growth superior to the wild type. Deleting *gacS* and *gacA* largely restored the phenotype exhibited by the evolved strains (Figure 11a, b), while the deletion of *fleQ* only partially restored the phenotype of the evolved strains (Figure 11c). Interestingly, *gacA* was also identified as a recurrent mutational target in an independent acetate-driven ALE experiment using a genome-reduced *P. putida* strain (Gurdo et al., 2026). The results thus confirmed that ALE-driven mutations in these genes disrupting their functions enhanced acetate utilization. Mutants of *gacS* and *gacA* were also found to exhibit significant fitness benefits during growth on acetate by Thompson et al. (2020).

To investigate how mutations affected the functions of PP_5292 and PP_1652, deletion- and overexpression mutants of both genes were created, and their growth behavior on acetate was investigated. The deletion of PP_5292, encoding CRC, restored the phenotype of the evolved strains, while overexpression did not improve growth compared to the wild type (Figure 11d). La Rosa et al. (2015) discovered that a mutant deficient in *crc* exhibits higher levels of ATP, NADH, and NADPH due to a rerouted metabolism and higher activity of ATP synthases. Since growth on acetate is assumed to require more energy than growth on glucose, the increased amount of available energy might cause the growth advantage of *P. putida* KT2440 Δ*crc* on acetate. Regarding *pfeS*-II, neither an overexpression nor a deletion of the gene was able to restore the phenotype of the evolved strains (Figure 11e). It would therefore be necessary to replicate the exact mutation that occurred in order to assign it a role in the improved phenotype on acetate.

Since the mutations cooccurred combined in the evolved strains and to test if combining beneficial mutations could further improve growth on acetate, *P. putida* KT2440 Δ*gacS* Δ*gacA* and *P. putida* KT2440 Δ*gacS* Δ*crc* were created. While *P. putida* KT2440 Δ*gacS* Δ*gacA* did not exhibit any advantage over the single gene KOs, the stacked mutant *P. putida* KT2440 Δ*gacS* Δ*crc* exhibited slightly improved growth compared to the single gene deletions (Figure 11 f). The evolved strains ALE and TALE exhibited growth rates of 0.27 h^-1^ and 0.28 h^-1^ and lag phases of one hour in this experiment, whereas the double deletion strain *P. putida* KT2440 Δ*gacS* Δ*crc* even exhibited a growth rate of 0.31 h^-1^ and a lag phase of one hour. In contrast, the wild type exhibited a growth rate of 0.25 h^-1^, but a lag phase of seven hours. Thus, reverse engineering restored the phenotype of the evolved strains and noticeably improved growth performance of *P. putida* KT2440. Since *gacS* and *gacA* are two parts of the same two-component system, deletion one of them is likely sufficient to disrupt the function, with no further advantage from deleting the second gene.

All in all, characterization of created deletion and overexpression mutants revealed that single deletions of either *gacS*, *gacA*, or *crc* are sufficient to improve growth of *P. putida* KT2440 on acetate to levels exhibited by strains generated by evolutionary engineering. The mutations found in evolved strains were thus confirmed to be causal for the presented phenotype. This finding is further supported by Gurdo et al. (2026), who linked attenuation of GacA-dependent regulation to reduced investment in costly cellular programs and improved conversion of acetate carbon into biomass. Nevertheless, the mutations might not be specific for growth on acetate. Mutations in *gacS* and *gacA* were also found during ALE of *P. putida* on glucose, ferulic acid, and coumaric acid (Mohamed et al. 2020), indicating that these adjustments are rather linked to the cultivation conditions than the applied carbon source. Further, none of the genes rationalized to be directly related to acetate metabolism (sections 3.4 and 3.5) was found to be affected by mutations during ALE experiments.

### 3.9 RNA-Seq data reveals altered energetic state of *P. putida* KT2440 TALE

The transcriptomes of the wild type during growth on glucose and on acetate, as well as the transcriptomes of *P. putida* KT2440 TALE during growth on acetate were analyzed to identify genes differentially expressed on acetate compared to glucose. The transcriptome of the evolved strain was additionally investigated to obtain further insights into differential gene expression and, thus, the effect of the mutations caused by ALE. For the wild type *P. putida* KT2440, 657 genes were differentially expressed during growth on acetate compared to growth on glucose, of which 440 genes were downregulated on acetate and 216 genes were upregulated (Figure S 3a). A similar range of differential expression was observed in *E. coli* by Oh et al. (2002), who found 354 genes to be upregulated and 370 genes to be downregulated upon growth on acetate. As expected, genes involved in sugar uptake and glycolytic pathways were significantly upregulated on glucose and, in turn, downregulated on acetate. Since the glyoxylate shunt is required during growth on acetate, the genes coding for isocitrate lyase and malate synthase G were upregulated, exhibiting a log2 fold change of 3.8 and 2.8, respectively, indicative of increases in the transcript level during growth on acetate. None of the acetate permease genes exhibited significant differential expression on acetate, again indicating that acetate does not enter the cell *via* one of the sodium/acetate symporters. In contrast, *acsA*-I (PP_4487), coding for an acetyl-CoA synthetase, was found to be overexpressed during growth on acetate (log2 fold change of 1.3), while no significant expression change was detected for *acs* (PP_3458). This result indicates that *acsA*-I was responsible for acetate activation and, thus, confirms results obtained with deletion and overexpression mutants in section 3.5. None of the genes targeted for reverse engineering were overexpressed during growth on acetate.

When gene expression of *P. putida* KT2440 TALE on acetate was compared to the wild type, 1,059 genes were found to be differentially expressed, with 852 of them downregulated and 207 upregulated (Figure S 3b). Notably, genes involved in the flagella synthesis were significantly downregulated (Table 6). Since flagella assembly and operation require high amounts of ATP (Martínez-García et al. 2014), downregulating the involved genes conserves energy. Of the 69 genes known to be involved in synthesis and operation of flagella, 41 were found to be downregulated in *P. putida* KT2440 TALE. None of these genes was found to be mutated in any of the evolved strains. However, the transcription factor FleQ, which is on top of the regulatory hierarchy regulating the transcription of flagella-related genes (Leal-Morales et al. 2022), was targeted by evolution in *P. putida* TALE and reverse engineering confirmed a loss of function mutation in the respective gene. By destruction of the function of FleQ, transcription of the genes regulated by this transcription factor is prevented. The evolved strain thus probably has an altered energy state and, therefore, can metabolize acetate more efficiently.

**Table 6:** Genes involved in flagella synthesis, which were found to be differentially expressed in *P. putida* KT2440 TALE compared to *P. putida* KT2440 during growth on acetate.

| Gene | Gene product | log2 fold change |
| --- | --- | --- |
| *flaG* | flagellar protein | -7.3 |
| *flgF* | flagellar basal-body rod protein | -5.4 |
| *flgB* | flagellar basal body rod protein | -5.1 |
| *flgG* | flagellar basal body rod protein | -5 |
| *flgC* | flagellar basal body rod protein | -4.9 |
| *fliE* | flagellar hook-basal body complex protein | -4.8 |
| *fliN* | flagellar motor switch protein | -4.7 |
| *flgK* | flagellar hook-associated protein | -4.6 |
| *fliD* | flagellar filament capping protein | -4.5 |
| *flgI* | flagellar basal body P-ring protein | -4.3 |
| *fliM* | flagellar motor switch protein | -4.3 |
| *flgA* | flagellar basal body P-ring formation chaperone | -4.2 |
| *flgD* | flagellar hook assembly protein | -4.2 |
| *flgH* | flagellar basal body L-ring protein | -4.1 |
| *fliS* | flagellar export chaperone | -4.1 |
| *flgJ* | flagellar assembly peptidoglycan hydrolase | -4.1 |
| *flgE* | flagellar hook protein | -3.9 |
| *fliL* | flagellar basal body-associated protein | -3.7 |
|  | flagellar hook-associated protein 3 | -3.7 |
| *fliO* | flagellar biosynthetic protein | -3.7 |
| *fliJ* | flagellar export protein | -3.5 |
| *fliP* | flagellar type III secretion system pore protein | -3.3 |
| *fliT* | flagellar assembly protein | -3.1 |
| *fliK* | flagellar hook-length control protein | -3.1 |
| *fliI* | flagellar protein export ATPase | -3 |
| *fliF* | flagellar basal-body MS-ring/collar protein | -2.8 |
|  | flagellar motor protein | -2.8 |
| *fliH* | flagellar assembly protein | -2.7 |
| *flhB* | flagellar biosynthesis protein | -2.3 |
| *motD* | flagellar motor protein | -2.3 |
| *fliG* | flagellar motor switch protein | -2.3 |
| *motB* | flagellar motor protein | -2 |
| *fleN* | flagellar synthesis regulator | -1.9 |
| *motA* | flagellar motor stator protein | -1.9 |
|  | flagellar brake protein | -1.8 |
| *flhA* | flagellar biosynthesis protein | -1.7 |
| *flhF* | flagellar biosynthesis protein | -1.6 |
| *fliQ* | flagellar biosynthesis protein | -1.6 |
| *flgN* | flagellar protein | -1.4 |
| *flgM* | flagellar biosynthesis anti-sigma factor | -1.3 |
| *fliK* | flagellar hook-length control protein | -1 |

Furthermore, the collected data were analyzed regarding the genes mutated during evolutionary engineering and considered for reverse engineering. While *gacS*, *crc*, and *fleQ* were not differentially expressed, *gacA* was drastically downregulated (log2 fold change of -6.8), and *pfeS*-II was slightly upregulated (log2 fold change of 1.0). The strong downregulation of *gacA* in *P. putida* KT2440 TALE aligns with the finding that deleting this gene improved growth on acetate, while overexpressing *pfeS*-II had only little impact. The RNA-seq data thus helped to confirm experimental results and provided insights into the altered metabolism of *P. putida* KT2440 TALE.

## 4 Discussion

*P. putida* is naturally able to metabolize a wide range of carbon sources and shows high tolerance towards toxic compounds (Poblete-Castro et al. 2012, Blanco-Romero et al. 2018). The tolerance of this strain towards acetate is higher than the tolerance of *E. coli* (Mutyala et al. 2023). When 0.5 g L^-1^ acetate was added to a culture of *E. coli*, the growth rate dropped by 48 % (Roe et al. 2002). Thus, acetate is considered a bearable substrate for *P. putida* KT2440 and was used for the bioproduction of PHA (Yang et al. 2019) and succinate (Mutyala et al. 2023) with *P. putida* already. Here, we investigated acetate metabolization in *P. putida* KT2440 using several approaches and tried to improve biomass and biosurfactant production by rational and evolutionary engineering.

### 4.1 Suitability of acetate as a carbon source for *P. putida*

Theoretical considerations imply that acetate is a suitable substrate for sustainable biotechnological processes and especially for acetyl-CoA-derived products, as the conversion of acetate to acetyl-CoA takes place with 100 % carbon efficiency. Using acetate derived from biotechnological CO_2_-balanced production or side streams can further contribute to a circular bioeconomy. However, compared to the conventional carbon source glucose, acetate exhibits a lower energy content. The results obtained in this study clearly demonstrate that acetate is a challenging carbon source. Although *P. putida* KT2440 is naturally able to metabolize acetate to biomass and secondary metabolites, the strain’s performance is negatively influenced by the C2 compound. This phenomenon can be partly explained by the fact that a portion of the supplied substrate is oxidized via the TCA cycle for energy generation, thereby meeting the elevated ATP demand associated with acetate activation to acetyl-CoA. As a result, the carbon is no longer available for biomass or product formation, and the energetic cost of substrate activation further constrains cellular energy, collectively explaining the observed phenotype. However, we also demonstrate that combining different approaches, including FBA, ALE with subsequent whole genome sequencing, as well as removing metabolic bottlenecks by genetic engineering, can improve the performance of *P. putida* KT2440, leaving much room for further improvement. Since acetate remains toxic, maintaining low acetate concentrations through process engineering should be considered an important strategy. Combined with strain engineering, this could lead to an economically viable process.

### 4.2 Adaptive laboratory evolution is an excellent tool for strain development

Adaptive laboratory evolution has been successfully used to improve growth of *P. putida* KT2440 on coumaric and ferulic acid (Mohamed et al. 2020), on xylose and galactose (Lim et al. 2021), on ethanol (Bator et al. 2020), and on syringyl aromatics (Mueller et al. 2022), among many others. By using ALE and subsequent whole genome sequencing, not only metabolic pathways but also regulatory mechanisms and tolerance mechanisms might be revealed.

Here, we demonstrated that growth of the strain on acetate could be improved by (T)ALE, resulting in evolved strains with higher growth rates and shorter lag phases. This work thus gives another example of ALE being a powerful tool to generate industrially relevant strains, which in some situations is more powerful than *a priori* rational strain engineering. Some genes that were found to be mutated in the here performed (T)ALE experiments have been shown to be the target of mutations in previous studies already (*gacS*, *uvrY* (also known as *gacA*), *fleQ*). Notably, *gacA* was also identified as a recurrent mutational target in an independent ALE experiment specifically selecting for improved acetate tolerance in *P. putida* (Gurdo et al., 2026). Mutations in all or some of these genes have been found after TALE on coumaric acid, but also after ALE on glucose (Mohamed et al. 2020), as well as after ALE on xylose (Lim et al. 2021), ethanol (Bator et al. 2020), or ionic liquids (Lim et al. 2020). Thus, the genes found to be affected by ALE are most likely not directly related to acetate metabolization. Since these genes were subject to mutations on various carbon sources, the results rather indicate mutations due to the cultivation conditions (minimal medium, stirred/shaken vessels) than the carbon source itself. GacS/GacA is involved in multiple cellular processes, including biofilm formation and motility (Song et al. 2023), traits that might not be needed in shaken suspended cultivation with abundant carbon source. A mutant deficient in *gacS* was found to exhibit reduced biofilm formation (Martínez-Gil et al. 2014). Mutations in this gene probably resulted in beneficial traits in the applied cultivation conditions. Mutations in *crc* leading to a disruption of the protein function probably resulted in a beneficial energetic state of the cell since deletion of *crc* was found to double the availability of ATP and NADPH in *P. putida* KT2440 on different carbon sources (La Rosa et al. 2015). Inactivation of FleQ, a transcriptional regulator involved in expression of flagellar genes, results in non-motile cells lacking flagella (Blanco-Romero et al. 2018). Since the assembly and operation of flagella is energy-intensive, inactivating *fleQ* relieves energy that can be used for other cellular processes. In this case, the surplus of energy might be used to cover the elevated ATP-demand computed by FBA (Ziegler et al. 2023), caused by the ATP-dependent activation of acetate, thus resulting in a growth advantage. Indeed, Zobel et al. (2017) observed an increase in biomass during growth on mixtures of glucose and formate, indicating that biomass yield on certain carbon sources in *P. putida* KT2440 is limited by energy availability. Several of the targets chosen for reverse engineering might be beneficial for growth on other substrates, too.

Although ALE thus helped improve the performance of *P. putida* KT2440 on acetate, it only shed limited light on acetate metabolization in this strain as many of the converged mutations were found to be not acetate specific. However, these results further legitimize a chassis cell that has been deprived of functions such as the flagellar transport as proposed, *e.g.*, by (Martínez-García et al. 2014).

### 4.3 Cross-membrane transport of acetate as weak organic acid

The transport of acetate across cellular membranes is a topic of discussion. Although it is commonly believed that acids can diffuse across membranes only in the unionized form and transport of dissociated acids relies on transport proteins, Axe and Bailey (1995) concluded that acetate can permeate the cell membrane in both, its anionic as well as its undissociated form.

In contrast, Kell (2021) ascribed a negligible role to diffusion in transport across biomembranes since these membranes consist not only of lipids but to a large extend of proteins, which prevent the formation of transient aqueous channels. Thus, cross-membrane transport of acetate was a subject of interest in this study. Three genes are annotated as acetate permeases in *P. putida* KT2440, thought to be involved in the transport of acetate across the membrane, but none of them was found to be essential for growth on acetate at the tested concentrations. In *P. fluorescence*, a deletion of *actP* did not cause a significant phenotype, either (Sepulveda and Lupas 2017), and it was previously shown that neither of the *actP* genes was upregulated during growth on acetate (Henriquez and Jung 2021), which was confirmed by RNA-seq data collected in this study, supporting this hypothesis. In the archaeon *Haloferax volcanii* growth on acetate was also unaffected in an *actP*-mutant, but acetate uptake rate was reduced drastically, which was not investigated here. However, the deletion of ActP was found to impair growth on acetate of *E. coli* (Bernal et al. 2016). Besides, *E. coli* has another transporter used for acetate uptake, the succinate-acetate transport protein SatP (Sá-Pessoa et al. 2013), of which no homolog was found in *P. putida* KT2440. Since a mutant of *E. coli* deficient in both, *actP* and *satP*, exhibited a strongly reduced acetate uptake rate, it has been concluded that *E. coli* takes up acetate *via* facilitated transport (Bernal et al. 2016).

One possible explanation for the lack of phenotype in the mutants could be acetate entering the cell *via* diffusion of the undissociated form. Given that only the uncharged form of acetate – being lipid-soluble – can enter the cell *via* diffusion, this is unlikely as this fraction is negligible. Since acetate has a pK_a_ of 4.75, at pH 7, 99.44 % of the molecules were calculated to be present in the ionic form using the Henderson-Hasselbalch equation. A more likely explanation is that other *P. putida* transporters compensate for the lack of acetate permeases. In *Corynebacterium glutamicum* a pyruvate transporter was shown to also mediate acetate transport (Jolkver et al. 2009), while in *E. coli* YaaH was shown to be involved in acetate uptake besides ActP (Sá-Pessoa et al. 2013). Thus, it is reasonable to assume that there is another, yet unidentified, transporter, or set of them, for acetate uptake into the cell in *P. putida* KT2440.

Despite its simple structure and well-characterized catabolism, acetate presents several challenges as a carbon source: its low energy content, the need for gluconeogenesis with its potentially reduced NADPH synthesis, and the stress associated with weak acid accumulation must all be considered. Realizing the potential of this promising carbon source requires a combined approach of metabolic and bioprocess engineering.

## 5 Authors’ contributions

**Melanie Filbig:** conceptualization, methodology, investigation, formal analysis, validation, visualization, data curation, writing – original draft, writing – review and editing. **Luisa Wachtendonk:** investigation. **Luca Hampe:** investigation. **Isabel Bator:** investigation. **Josefin Johnsen:** investigation, methodology. **Elsayed T. Mohamed:** investigation, methodology. **Nicolás Gurdo:** investigation, methodology. **Johannes Parschau1:** investigation. **Pablo I. Nikel:** resources, supervision. **Adam M. Feist:** methodology, formal analysis, resources, supervision, writing – review and editing. **Till Tiso:** conceptualization, methodology, formal analysis, resources, supervision, funding acquisition, project administration, writing – review and editing. **Lars M. Blank:** conceptualization, methodology, formal analysis, resources, supervision, funding acquisition, project administration, writing – review and editing.

## 6 Acknowledgments

Authors M.F., T.T., and L.M.B. were partially supported by the Deutsche Forschungsgemeinschaft (DFG, German Research Foundation) under Germany’s Excellence Strategy – Exzellenzcluster 2186 “The Fuel Science Center” ID: 390919832. T.T. and L.M.B. acknowledge funding from the European Union’s Horizon 2020 research and innovation program under grant agreement no. 870294 for the project MIX-UP. T.T. was partially funded by the Bioeconomy Science Center (BioSC), which is financially supported by the Ministry of Culture and Science within the framework of the NRW Strategieprojekt BioSC (No. 313/323-400-00213). Funding for this work was also provided by the Novo Nordisk Foundation under the Grant number NNF24SA0100980. This material is based upon work at the Joint BioEnergy Institute (JBEI) supported by the U.S. Department of Energy, Office of Science, Biological and Environmental Research Program under contract DE-AC02-05CH11231 with Lawrence Berkeley National Laboratory.

