## Supplementary Information for "Improving acetate metabolism of *P. putida* KT2440 by evolutionary and rational engineering"

To allow for microbial growth on acetate, generally, three steps are required: (I) acetate entering the cell, (II) activation of acetate into acetyl-CoA, and (III) assimilation of acetate *via* the TCA and glyoxylate cycle ().

The first crucial step for acetate metabolization is how acetate enters the cell. Acetate can either enter the cell by diffusion in its undissociated form, which is lipid-soluble, simply driven by a concentration gradient or by active transport of the dissociated, lipid-insoluble, acetate molecules across the membrane using $H^+$ /monocarboxylic acid symporters (PMCT) or sodium/solute symporters as acetate permeases (ActP). When in the cell, two ATP-dependent pathways exist for the activation of acetate to acetyl-CoA: the acetyl-CoA synthetase (ACS) pathway and the AckA-PTA pathway. While acetyl-CoA synthetase irreversibly converts acetate to acetyl-CoA under the consumption of ATP, AckA-PTA involves a reversible two-step path: AckA catalyzes the formation of acetyl phosphate from acetate followed by subsequent conversion to acetyl-CoA by PTA (Kutscha and Pflügl 2020). The ACS pathway requires more ATP than the AckA-PTA pathway, but ACS has the advantage of a 35-fold higher affinity to acetate, rendering it the more effective mechanism for acetate activation (Wolfe 2005, Kiefer et al. 2021).

Acetyl-CoA generated from acetate in a next step enters tricarboxylic acid cycle (TCA) or glyoxylate shunt (Kiefer et al. 2021). The glyoxylate shunt circumvents $CO_2$-producing reactions in the TCA cycle and thus enables the formation of C4 molecules from C2 substrates as acetate. Oxaloacetate generated in the TCA cycle serves as a precursor for amino acid biosynthesis, as well as gluconeogenic reactions. Gluconeogenic reactions are required during growth on substrates entering the central carbon metabolism at the stage of acetyl-CoA to supply sugar phosphates.

**Table S 1: Oligonucleotides used in this work.**

| Primer | Product | Template | Orientation | Sequence |
|---|---|---|---|---|
| BW13 | pEMG_mcs | pEMG | fwd | TTTGCACTGCCGGTAGAAC |
| BW14 | pEMG_mcs | pEMG | rev | AATACGCAAACCGCCTCTC |

| | | | | |
|---|---|---|---|---|
| 4_Pput-glmS | Mapping *glmS* | *att*Tn7 | fd | AGTCAGAGTTACGGAATTGTAGG |
| 5_Pput-glmS | Mapping *glmS* | *att*Tn7 | rev | GTCGAGAAAATTGCCGAGCT |
| 43_seq | pBGxx msc | pBGxx | fwd | ATCAAACATCGACCCACGGCGTAAC |
| 62 | pBGxx msc | pBGxx | rev | TAGTCGCCAGGGTTTTCC |
| MF68 | TS1 PP_1743 (*actP*-I) | *P. putida* KT2440 | fwd | agggataacagggtaatctgGCCAATGTTGTTTCTTCGCG |
| MF69 | TS1 PP_1743 (*actP*-I) | *P. putida* KT2440 | rev | gggattgatgTGTTGCGCCTCCTTCAGG |
| MF70 | TS2 PP_1743 (*actP*-I) | *P. putida* KT2440 | fwd | aggcgcaacaCATCAATCCCCTGCATTG |
| MF71 | TS2 PP_1743 (*actP*-I) | *P. putida* KT2440 | rev | ccgggtaccgagctcgaattGAATCTTTACCAACCCGC |
| MF72 | Mapping PP_1743 (*actP*-I) | *P. putida* KT2440 | fwd | GATGCTGGCGATCCAGGAAG |
| MF73 | Mapping PP_1743 (*actP*-I) | *P. putida* KT2440 | rev | CGGGCATGATTGGGTAGACG |
| MF74 | TS1 PP_2797 (*actP*-II) | *P. putida* KT2440 | fwd | agggataacagggtaatctgTTCAAGGCGCCGCGCCAT |
| MF75 | TS1 PP_2797 (*actP*-II) | *P. putida* KT2440 | rev | cgcgggtaacGCGCCCTGCTCCTCTTCC |
| MF76 | TS2 PP_2797 (*actP*-II) | *P. putida* KT2440 | fwd | agcagggcgcGTTACCCGCGGCCGCAAG |
| MF77 | TS2 PP_2797 (*actP*-II) | *P. putida* KT2440 | rev | ccgggtaccgagctcgaattTCGAGGAAGCCGTACGCC |
| MF78 | Mapping PP_2797 (*actP*-II) | *P. putida* KT2440 | fwd | ACACCTCACGCTTTCGTCAC |
| MF79 | Mapping PP_2797 (*actP*-II) | *P. putida* KT2440 | rev | TGTTTCGAAGACGGCAAGGC |
| MF80 | TS1 PP_3272 (*actP*-III) | *P. putida* KT2440 | fwd | agggataacagggtaatctgGCCACCCTGGAAGCTGGA |
| MF81 | TS1 PP_3272 (*actP*-III) | *P. putida* KT2440 | rev | agtgcgcacaCTTCAAGGACGAACAGACATGTCG |
| MF82 | TS2 PP_3272 (*actP*-III) | *P. putida* KT2440 | fwd | gtccttgaagTGTGCGCACTCCTGCTTG |
| MF83 | TS2 PP_3272 (*actP*-III) | *P. putida* KT2440 | rev | ccgggtaccgagctcgaattTGCCAGACCTACCCGGTG |
| MF84 | Mapping PP_3272 (*actP*-III) | *P. putida* KT2440 | fwd | AGTCACGCAGGTTGGCCTTG |
| MF85 | Mapping PP_3272 (*actP*-III) | *P. putida* KT2440 | rev | GGCGAAGTGGAGATGGACAG |

| | | | | |
|---|---|---|---|---|
| MF270 | TS1 PP_4946 (*putP*) | *P. putida* KT2440 | fwd | agggataacagggtaatctgTTTTCATGTTTTCCTTGAGGAGC |
| MF271 | TS1 PP_4946 (*putP*) | *P. putida* KT2440 | rev | ggagcaagaaCACCCGTTGAATTGGCGG |
| MF272 | TS2 PP_4946 (*putP*) | *P. putida* KT2440 | fwd | tcaacgggtgTTCTTGCTCCTTTCGGGAGCAAAAAACCG |
| MF273 | TS2 PP_4946 (*putP*) | *P. putida* KT2440 | rev | atccccgggtaccgagctcgGGCGGTAGGCGGCGGTGA |
| MF278 | Mapping PP_4946 (*putP*) | *P. putida* KT2440 | fwd | TCGCCTTGCAGGTCGTACAG |
| MF279 | Mapping PP_4946 (*putP*) | *P. putida* KT2440 | rev | AGCTGCTTTGAGTCGCTCAC |
| MF274 | TS1 PP_4524 (sodium-solute symporter) | *P. putida* KT2440 | fwd | agggataacagggtaatctgCCGACGACTTCAACTGGAG |
| MF275 | TS1 PP_4524 (sodium-solute symporter) | *P. putida* KT2440 | rev | ggtacagcagTGGGGGTCTCCCGATTATC |
| MF276 | TS2 PP_4524 (sodium-solute symporter) | *P. putida* KT2440 | fwd | gagacccccaCTGCTGTACCGGCCCTTTC |
| MF277 | TS2 PP_4524 (sodium-solute symporter) | *P. putida* KT2440 | rev | atccccgggtaccgagctcgTCAGGCGGTCACCTGGCT |
| MF280 | Mapping PP_4524 (sodium-solute symporter) | *P. putida* KT2440 | fwd | AGATCATTCGCGGCTGCCAC |
| MF281 | Mapping PP_4524 (sodium-solute symporter) | *P. putida* KT2440 | rev | AGGATTGATTTGCCGTGAGG |
| MF192 | TS1 PP_3458 (*acs*) | *P. putida* KT2440 | fwd | agggataacagggtaatctgGAGCGACGACCTGGTGATTTC |
| MF193 | TS1 PP_3458 (*acs*) | *P. putida* KT2440 | rev | gagaacgcccGGTTCACCCGCGAAAGGG |
| MF194 | TS2 PP_3458 (*acs*) | *P. putida* KT2440 | fwd | cgggtgaaccGGGCGTTCTCTTGCTGTTC |
| MF195 | TS2 PP_3458 (*acs*) | *P. putida* KT2440 | rev | atccccgggtaccgagctcgACCGCTGTATGGCTACCTG |
| MF196 | Mapping PP_3458 (*acs*) | *P. putida* KT2440 | fwd | CCTGCTCCATCCTCATTC |
| MF197 | Mapping PP_3458 (*acs*) | *P. putida* KT2440 | rev | GGCCAAGTACGACATCAAAC |
| MF204 | TS1 PP_3724 (acyl-CoA synthetase) | *P. putida* KT2440 | fwd | agggataacagggtaatctgGCGTACCACCTCACCGCT |
| MF205 | TS1 PP_3724 (acyl-CoA synthetase) | *P. putida* KT2440 | rev | gagaacactcCCTTACCCTGACAGCCGG |

| | | | | |
|---|---|---|---|---|
| MF206 | TS2 PP_3724 (acyl-CoA synthetase) | *P. putida* KT2440 | fwd | cagggtaaggGAGTGTTCTCCTGGCGCTC |
| MF207 | TS2 PP_3724 (acyl-CoA synthetase) | *P. putida* KT2440 | rev | atccccgggtaccgagctcgCGTGGTTATCACACCT ACGAG |
| MF208 | Mapping PP_3724 (acyl-CoA synthetase) | *P. putida* KT2440 | fwd | GTACTGAGTTGCTGCAACTG |
| MF209 | Mapping PP_3724 (acyl-CoA synthetase) | *P. putida* KT2440 | rev | CCGACACCTACGAGCACAAC |
| MF210 | TS1 PP_4487 (*acsA*-I) | *P. putida* KT2440 | fwd | agggataacagggtaatctgGGTAACAGCTGCCCGA TATG |
| MF211 | TS1 PP_4487 (*acsA*-I) | *P. putida* KT2440 | rev | ggtaacacagACCCTTCCCGGCAATACC |
| MF212 | TS2 PP_4487 (*acsA*-I) | *P. putida* KT2440 | fwd | cgggaagggtCTGTGTTACCTCGGTGTAATAG |
| MF213 | TS2 PP_4487 (*acsA*-I) | *P. putida* KT2440 | rev | atccccgggtaccgagctcgAGTAACTGTACTGTTC AACC |
| MF214 | Mapping PP_4487 (*acsA*-I) | *P. putida* KT2440 | rev | CGCGGTATTCGAGAATGTGC |
| MF215 | Mapping PP_4487 (*acsA*-I) | *P. putida* KT2440 | fwd | TTTCAGCAGCCTGCGACAC |
| MF216 | TS1 PP_4702 (*acsA*-II) | *P. putida* KT2440 | fwd | agggataacagggtaatctgAAACCCGCGAGGAAG CCG |
| MF217 | TS1 PP_4702 (*acsA*-II) | *P. putida* KT2440 | rev | gccagatttgCGATGGACTGCAACGGCTATTC |
| MF218 | TS2 PP_4702 (*acsA*-II) | *P. putida* KT2440 | fwd | cagtccatcgCAAATCTGGCCGCCCTGTAC |
| MF219 | TS2 PP_4702 (*acsA*-II) | *P. putida* KT2440 | rev | atccccgggtaccgagctcgGGTTGCGGTAACGCAG GTG |
| MF220 | Mapping PP_4702 (*acsA*-II) | *P. putida* KT2440 | fwd | TTGATTCCGGCCGATTTC |
| MF221 | Mapping PP_4702 (*acsA*-II) | *P. putida* KT2440 | rev | GGCGTATGTGTTGCAGATAG |
| MF226 | pEMG-PP_0340_BG42 | pEMG-PP_0340_BG42 | fwd | ttcggcgtaaGAATTCGAGCTCGGTACC |
| MF227 | pEMG-PP_0340_BG42 | pEMG-PP_0340_BG42 | rev | gcgcgggcatTAGAAAACCTCCTTAGCATG |
| MF228 | PP_4487 (*acsA*-I) | *P. putida* KT2440 | fwd | aggttttctaATGCCCGCGCCAGAGCGT |

| | | | | |
|---|---|---|---|---|
| MF229 | PP_4487 (*acsA*-I) | *P. putida* KT2440 | rev | gctcgaattcTTACGCCGAAGCCAGGTTCATCGC |
| MF242 | Mapping PP_0340 | *P. putida* KT2440 | fwd | AAGCCTGGACCTGGGAACAC |
| MF243 | Mapping PP_0340 | *P. putida* KT2440 | rev | GCGCTTCGATCTGCAGGTTG |
| MF236 | pBGffg_lin | $pBG14_{ffg}$-msf*gfp* | fwd | GAATTCGAGCTCGGTACC |
| MF237 | pBGffg_lin | $pBG14_{ffg}$-msf*gfp* | rev | TAGAAAACCTCCTTAGCATG |
| MF238 | PP_4116 (*aceA*) | *P. putida* KT2440 | fwd | catgctaaggaggttttctaATGGCACTGACACGCGAAC |
| MF239 | PP_4116 (*aceA*) | *P. putida* KT2440 | rev | atccagtcatTCAGTGGAACTGCTCTTCTTC |
| MF240 | PP_0356 (*glcB*) | *P. putida* KT2440 | fwd | gttccactgaATGACTGGATACGTTCAAGTC |
| MF241 | PP_0356 (*glcB*) | *P. putida* KT2440 | rev | cgggtaccgagctcgaattcTTACAACCCGTTACGCGC |
| MF332 | TS1 PP_1650 (*gacS*) | *P. putida* KT2440 | fwd | agggataacagggtaatctgGGTCGAAGCCGGTCAGCC |
| MF333 | TS1 PP_1650 (*gacS*) | *P. putida* KT2440 | rev | ggaggcgagtGGGGATCTGGGCCACGCT |
| MF334 | TS2 PP_1650 (*gacS*) | *P. putida* KT2440 | fwd | ccagatccccACTCGCCTCCTCTGATGTGCCTG |
| MF335 | TS2 PP_1650 (*gacS*) | *P. putida* KT2440 | rev | atccccgggtaccgagctcgCGGCGCCGCGCTGAAAGC |
| MF336 | Mapping PP_1650 (*gacS*) | *P. putida* KT2440 | fwd | AGGCTGATGATGCGGGCTTC |
| MF337 | Mapping PP_1650 (*gacS*) | *P. putida* KT2440 | rev | TGTCCAGCAGGCGGTATTCG |
| MF338 | TS1 PP_4099 (*uvrY*/*gacA*) | *P. putida* KT2440 | fwd | agggataacagggtaatctgGGCCGGGTGCGGTTGGCG |
| MF339 | TS1 PP_4099 (*uvrY*/*gacA*) | *P. putida* KT2440 | rev | aggtgtgtgcAACCTCATGTCCCAAGTCTTTGATGCCAGCG |
| MF340 | TS2 PP_4099 (*uvrY*/*gacA*) | *P. putida* KT2440 | fwd | acatgaggttGCACACACCTCGTCAGCAGG |
| MF341 | TS2 PP_4099 (*uvrY*/*gacA*) | *P. putida* KT2440 | rev | atccccgggtaccgagctcgGGCGGGCAGCCCGTACTT |
| MF342 | Mapping PP_4099 (*uvrY*/*gacA*) | *P. putida* KT2440 | fwd | TCCATCTCGGCGTTCAACTC |

| | | | | |
|---|---|---|---|---|
| MF343 | Mapping PP_4099 (*uvrY*/*gacA*) | *P. putida* KT2440 | rev | TGTACCAGCGTCATGGCTTC |
| MF344 | TS1 PP_1652 (*pfeS*-II) | *P. putida* KT2440 | fwd | agggataacagggtaatctgGCAGGCACAGCACCACAC |
| MF345 | TS1 PP_1652 (*pfeS*-II) | *P. putida* KT2440 | rev | ccttcaagtcGGTTCACTCCGCCTCGCT |
| MF346 | TS2 PP_1652 (*pfeS*-II) | *P. putida* KT2440 | fwd | ggagtgaaccGACTTGAAGGGAGGAGGAG |
| MF347 | TS2 PP_1652 (*pfeS*-II) | *P. putida* KT2440 | rev | atccccgggtaccgagctcgAAGCGGCATCATGGGCATC |
| MF348 | Mapping PP_1652 (*pfeS*-II) | *P. putida* KT2440 | fwd | CTGCCGCACCATTGCATCTC |
| MF349 | Mapping PP_1652 (*pfeS*-II) | *P. putida* KT2440 | rev | CTATGCGTTCGAGCACTTCC |
| MF350 | TS1 PP_5292 (CRC protein) | *P. putida* KT2440 | fwd | agggataacagggtaatctgGATGATCTGCATGACCTC |
| MF351 | TS1 PP_5292 (CRC protein) | *P. putida* KT2440 | rev | caatggccttAAATGGCCCCATAAATCTC |
| MF352 | TS2 PP_5292 (CRC protein) | *P. putida* KT2440 | fwd | ggggccatttAAGGCCATTGGGGCTGCA |
| MF353 | TS2 PP_5292 (CRC protein) | *P. putida* KT2440 | rev | atccccgggtaccgagctcgCAACGCCATGCTCGCTTTG |
| MF354 | Mapping PP_5292 (CRC protein) | *P. putida* KT2440 | fwd | CAGCCGGCAGATGCTGTTTC |
| MF355 | Mapping PP_5292 (CRC protein) | *P. putida* KT2440 | rev | AAGACTCCGCAGCCGAAACC |
| MF378 | PP_1650 (*gacS*) | *P. putida* KT2440 | fwd | catgctaaggaggttttctaGTGCTCGATCGCTTGGGAATC |
| MF379 | PP_1650 (*gacS*) | *P. putida* KT2440 | rev | cgggtaccgagctcgaattcTCAAGCGCTCAATCTGGC |
| MF380 | PP_1652 (*pfeS*-II) | *P. putida* KT2440 | fwd | catgctaaggaggttttctaATGCTCGACCGCCATTCG |
| MF381 | PP_1652 (*pfeS*-II) | *P. putida* KT2440 | rev | cgggtaccgagctcgaattcCTACCATACAAGCGGGTTTACC |
| MF382 | PP_4099 (*uvrY*/*gacA*) | *P. putida* KT2440 | fwd | catgctaaggaggttttctaTTGATTAGGGTCTTAGTGGTC |
| MF383 | PP_4099 (*uvrY*/*gacA*) | *P. putida* KT2440 | rev | cgggtaccgagctcgaattcTTACAGGCTTGCGTCAAC |
| MF384 | PP_5292 (CRC protein) | *P. putida* KT2440 | fwd | catgctaaggaggttttctaATGCGGATCATCAGTGTG |

| | | | | |
|---|---|---|---|---|
| MF385 | PP_5292 (CRC protein) | *P. putida* KT2440 | rev | cgggtaccgagctcgaattcTTAGATGGTCAGCGTCCAG |
| IB7 | Mapping PP_4219-PP_4221 (pvd) | *P. putida* KT2440 | fwd | ACCCATACGCATGAAGTC |
| IB8 | Mapping PP_4219-PP_4221 (pvd) | *P. putida* KT2440 | rev | TACTGCTGCGTGGTTTCG |
| IB13 | Mapping PP_1277-PP_1288 (alg) | *P. putida* KT2440 | fwd | TCTTGCCAGACCACGAAC |
| IB14 | Mapping PP_1277-PP_1288 (alg) | *P. putida* KT2440 | rev | TACTACAGTGCCGAGCAG |
| IB19 | Mapping PP_4328-PP_4344 (flag1) | *P. putida* KT2440 | fwd | ATGGCGAAGAACACCAAC |
| IB20 | Mapping PP_4328-PP_4344 (flag1) | *P. putida* KT2440 | rev | TCCACCGAGTCATGAAGG |
| IB25 | Mapping PP_4351-PP_4397 (flag2) | *P. putida* KT2440 | fwd | AAACGGGATGGCACAAGC |
| IB26 | Mapping PP_4351-PP_4397 (flag2) | *P. putida* KT2440 | rev | GAGCCGAAGTTCTTCATC |
| IB47 | Mapping PP_2634-PP_2638 (bcs) | *P. putida* KT2440 | fwd | AGTGACCTGGATGTCTTG |
| IB48 | Mapping PP_2634-PP_2638 (bcs) | *P. putida* KT2440 | rev | TCACCGCCACAGTCTTTC |
| IB53 | Mapping PP_3132-PP_3142 (pea) | *P. putida* KT2440 | fwd | CGTAACCCAGTGCAATCG |
| IB54 | Mapping PP_3132-PP_3142 (pea) | *P. putida* KT2440 | rev | ATGCGCCAACTGGAAGAG |
| IB59 | Mapping PP_1795-PP_1788 (peb) | *P. putida* KT2440 | fwd | GCGCCATAATCAATGCTG |
| IB60 | Mapping PP_3132-PP_3142 (pea) | *P. putida* KT2440 | rev | TACACATCCCTGCTCAAC |
| IB79 | Mapping PP_0168 (*lapA*) | *P. putida* KT2440 | fwd | CCTTGAATCGGTGTTGAG |
| IB80 | Mapping PP_0168 (*lapA*) | *P. putida* KT2440 | rev | GTCCAGGCCTAAGATCTC |

| | | | | |
|---|---|---|---|---|
| IB85 | Mapping PP_0806 (*lapF*) | *P. putida* KT2440 | fwd | GTACTGCGACTGGTACTC |
| IB86 | Mapping PP_0806 (*lapF*) | *P. putida* KT2440 | rev | CTGTTCCTCGACGAAGTG |
| IB61 | Mapping PP_5003-PP_5008 (*phaCZCDFI*) | *P. putida* KT2440 | fwd | AGCGTTTGCTCGAAGAAGTG |
| IB154 | Mapping PP_5003-PP_5008 (*phaCZCDFI*) | *P. putida* KT2440 | rev | GCAATAGATCCGGTAGGG |

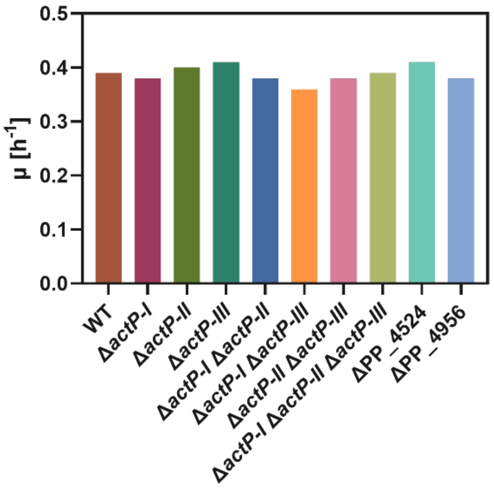


**Figure S 1: Growth rates of mutants of *P. putida* KT2440 on 5 g L$^{-1}$ acetate determined using the Growth Profiler (EnzyScreen BV, Heemstede, the Netherlands).**

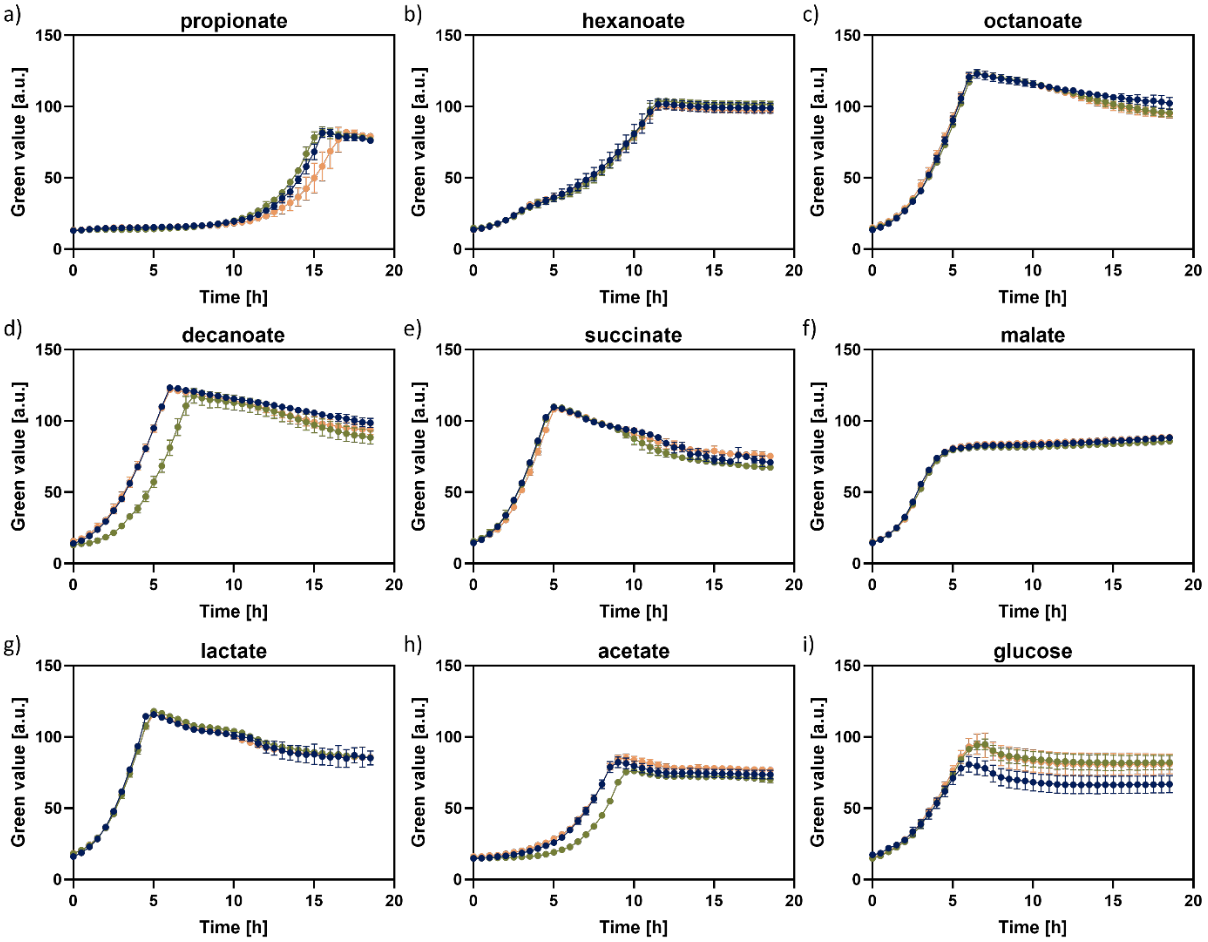


**Figure S 2: Biomass formation of *P. putida* KT2440 (blue), *P. putida* KT2440 Δ*actP*I-III (green) and *P. putida* KT2440 Δ*actP*I-III ΔPP_4524 ΔPP_4946 (orange) on 0.12 Cmol of different organic acids compared to glucose.** a) propionate, b) hexanoate, c) octanoate, d) decanoate, e) succinate, f) malate, g) lactate, h) acetate and i) glucose. Cultivation was performed in 24-well microtiter plates with transparent bottom with 1.5 ml filling volume in a growth profiler. Green value was determined as a measure of culture turbidity, *i.e.*, biomass formation. Error bars represent the standard deviation of three biological replicates.

**Table S 2: All mutations found in ALE experiments performed in this study.**

| Position | Mutation Type | Sequence Change | Gene | Product | Details | Experiment |
|---|---|---|---|---|---|---|
| 6,547 | SNP | T→C | yidC | membrane protein insertase YidC | N536S (AAC→AGC) | KT2440 GR18 ALE |
| 17,200 | INS | (G)6→7 | PP_RS00070 | hypothetical protein | coding (612/639 nt) | KT2440 GR18 ALE |
| 27,573 | INS | (A)9→10 | PP_RS00120, PP_RS00125 | hypothetical protein/phosphoethanolamine--lipid A transferase | intergenic (+118/+277) | KT2440 GR18 ALE |
| 167,974 | SNP | C→T | PP_RS00810 | LysR family transcriptional regulator | Q90Q (CAG→CAA) | KT2440 GR18 ALE |
| 234,926 | SNP | A→G | PP_RS00980 | glutathione S-transferase | I96V (ATC→GTC) | KT2440 GR18 ALE |
| 268,750 | INS | (G)6→7 | PP_RS01150, desA | sensor domain-containing diguanylate cyclase/delta-9 fatty acid desaturase DesA | intergenic (-81/+19) | KT2440 GR18 ALE |
| 293,916 | SNP | T→C | ssuC | aliphatic sulfonate ABC transporter permease SsuC | R38R (CGT→CGC) | KT2440 GR18 ALE |
| 313,169 | SNP | A→G | fdhD | formate dehydrogenase accessory sulfurtransferase FdhD | S15P (TCC→CCC) | KT2440 GR18 ALE |
| 323,387 | SNP | C→T | PP_RS01405 | TonB-dependent siderophore receptor | R285H (CGT→CAT) | KT2440 GR18 ALE |
| 325,039 | SNP | G→T | PP_RS01410 | OprD family porin | E109* (GAA→TAA) | KT2440 TALE |
| 325,289 | DEL | (AGCGCCG)2→1 | PP_RS01410 | OprD family porin | coding (575-581/1320 nt) | KT2440 TALE |
| 325,374 | SNP | C→G | PP_RS01410 | OprD family porin | Y220* (TAC→TAG) | KT2440 TALE |
| 450,940 | DEL | Δ53 bp | [PP_RS01965] | [PP_RS01965] | | KT2440 GR18a ALE |
| 451,028 | SNP | A→G | PP_RS01965 | LysR family transcriptional regulator | V282A (GTG→GCG) | KT2440 GR18a ALE |
| 451,146 | SNP | G→T | PP_RS01965 | LysR family transcriptional regulator | R243S (CGC→AGC) | KT2440 GR18 ALE |
| 451,614 | SNP | C→A | PP_RS01965 | LysR family transcriptional regulator | G87C (GGC→TGC) | KT2440 GR18a ALE |
| 453,404 | SNP | C→T | PP_RS01970, PP_RS01975 | aminotransferase class III-fold pyridoxal phosphate-dependent enzyme/YqaE/Pmp3 family membrane protein | intergenic (+41/-107) | KT2440 GR18 ALE |

| | | | | | | |
|---|---|---|---|---|---|---|
| 453,405 | SNP | G→C | PP_RS01970, PP_RS01975 | aminotransferase class III-fold pyridoxal phosphate-dependent enzyme/YqaE/Pmp3 family membrane protein | intergenic (+42/-106) | KT2440 GR18 TALE |
| 472,652 | SNP | G→A | rpoD | RNA polymerase sigma factor RpoD | R95C (CGC→TGC) | KT2440 ALE |
| 472,781 | SNP | C→T | rpoD | RNA polymerase sigma factor RpoD | D52N (GAC→AAC) | KT2440 ALE |
| 521,778 | SNP | G→A | PP_RS02285 | peptidoglycan DD-metalloendopeptidase family protein | R315C (CGC→TGC) | KT2440 GR18 TALE |
| 539,585 | SNP | G→T | rpoB | DNA-directed RNA polymerase subunit beta | V655F (GTC→TTC) | KT2440 GR18 TALE |
| 541,960 | SNP | A→G | rpoC | DNA-directed RNA polymerase subunit beta' | D67G (GAC→GGC) | KT2440 TALE |
| 545,079 | SNP | G→T | rpoC | DNA-directed RNA polymerase subunit beta' | V1107F (GTC→TTC) | KT2440 GR18 TALE |
| 545,395 | SNP | A→G | rpoC | DNA-directed RNA polymerase subunit beta' | D1212G (GAC→GGC) | KT2440 GR18 TALE |
| 699,722 | SNP | A→G | PP_RS03155, PP_RS03160 | tRNA-Ala/23S ribosomal RNA | intergenic (+115/-129) | KT2440 TALE |
| 699,722 | SNP | A→G | PP_RS03155, PP_RS03160 | tRNA-Ala/23S ribosomal RNA | intergenic (+115/-129) | KT2440 ALE |
| 699,722 | SNP | A→G | PP_RS03155, PP_RS03160 | tRNA-Ala/23S ribosomal RNA | intergenic (+115/-129) | KT2440 GR18 ALE |
| 699,722 | SNP | A→G | PP_RS03155, PP_RS03160 | tRNA-Ala/23S ribosomal RNA | intergenic (+115/-129) | KT2440 GR18 TALE |
| 699,722 | SNP | A→G | PP_RS03155, PP_RS03160 | tRNA-Ala/23S ribosomal RNA | intergenic (+115/-129) | KT2440 GR18a ALE |
| 721,559 | SNP | C→T | PP_RS03260 | ABC transporter ATP-binding protein | G147S (GGC→AGC) | KT2440 GR18 ALE |
| 744,901 | AMP | 2,100 bp x 2 | [PP_RS03405],PP_RS03410,tnpB | [PP_RS03405],PP_RS03410,tnpB | duplication | KT2440 GR18 TALE |
| 782,971 | SNP | C→T | PP_RS03595 | EAL domain-containing protein | M985I (ATG→ATA) | KT2440 TALE |
| 810,683 | SNP | C→T | rimI | ribosomal protein S18-alanine N-acetyltransferase | G126G (GGC→GGT) | KT2440 TALE |
| 836,643 | SNP | A→G | ychF | redox-regulated ATPase YchF | V238A (GTG→GCG) | KT2440 GR18 ALE |

| | | | | | | |
|---|---|---|---|---|---|---|
| 974,551 | DEL | (T)9→8 | secF, PP_RS04435 | protein translocase subunit SecF/glycine zipper 2TM domain-containing protein | intergenic (+61/-58) | KT2440 GR18 ALE |
| 976,596 | SNP | C→T | trmJ | tRNA (cytosine(32)/uridine(32)-2'-O)-methyltransferase TrmJ | H103Y (CAT→TAT) | KT2440 GR18 ALE |
| 1,162,637 | SNP | C→T | PP_RS05325 | carbohydrate porin | G135G (GGC→GGT) | KT2440 GR18 ALE |
| 1,164,953 | SNP | A→G | hexR | DNA-binding transcriptional regulator HexR | F151S (TTC→TCC) | KT2440 GR18 ALE |
| 1,186,063 | SNP | T→C | purL | phosphoribosylformylglycinamidine synthase | V423A (GTG→GCG) | KT2440 GR18 ALE |
| 1,302,915 | DEL | (AGC)4→3 | PP_RS05915 | ABC transporter ATP-binding protein | coding (459-461/702 nt) | KT2440 GR18 TALE |
| 1,303,453 | SNP | T→G | livG | high-affinity branched-chain amino acid ABC transporter ATP-binding protein LivG | T231P (ACC→CCC) | KT2440 TALE |
| 1,304,424 | DEL | (CGGCGAAGC)2→1 | PP_RS05925 | high-affinity branched-chain amino acid ABC transporter permease LivM | coding (965-973/1257 nt) | KT2440 TALE |
| 1,305,193 | DEL | Δ7 bp | PP_RS05925 | high-affinity branched-chain amino acid ABC transporter permease LivM | coding (198-204/1257 nt) | KT2440 TALE |
| 1,305,770 | DEL | (AGGCGCGGC)2→1 | livH | high-affinity branched-chain amino acid ABC transporter permease LivH | coding (539-547/924 nt) | KT2440 TALE |
| 1,306,946 | DEL | Δ2 bp | PP_RS05935 | branched-chain amino acid ABC transporter substrate-binding protein | coding (671-672/1116 nt) | KT2440 TALE |
| 1,307,095 | SNP | C→T | PP_RS05935 | branched-chain amino acid ABC transporter substrate-binding protein | E175K (GAA→AAA) | KT2440 GR18 TALE |
| 1,309,196 | SNP | A→G | PP_RS05945 | NAD(P)-dependent oxidoreductase | H239R (CAC→CGC) | KT2440 GR18a ALE |
| 1,375,237 | SNP | C→T | PP_RS06210 | potassium transporter Kup | A286V (GCC→GTC) | KT2440 GR18 ALE |
| 1,386,816 | SNP | C→T | PP_RS06255, PP_RS06260 | OprD family porin/HIT domain-containing protein | intergenic (-360/-331) | KT2440 GR18 TALE |
| 1,390,075 | SNP | T→C | PP_RS06285 | zinc ribbon domain-containing protein | L66P (CTG→CCG) | KT2440 GR18 ALE |
| 1,398,106 | SNP | C→T | tolA | cell envelope integrity protein TolA | P352S (CCG→TCG) | KT2440 GR18 ALE |

| | | | | | | |
|---|---|---|---|---|---|---|
| 1,398,565 | SNP | C→T | tolB | Tol-Pal system beta propeller repeat protein TolB | A123V (GCG→GTG) | KT2440 GR18 ALE |
| 1,631,753 | SNP | T→C | lepA | translation elongation factor 4 | G169G (GGT→GGC) | KT2440 GR18 ALE |
| 1,764,135 | SNP | C→T | PP_RS08105, PP_RS28520 | DUF3168 domain-containing protein/phage tail protein | intergenic (+108/-33) | KT2440 ALE |
| 1,802,517 | SNP | T→C | dnaE | DNA polymerase III subunit alpha | V834A (GTC→GCC) | KT2440 GR18 ALE |
| 1,802,558 | SNP | G→T | dnaE | DNA polymerase III subunit alpha | V848L (GTG→TTG) | KT2440 GR18 TALE |
| 1,823,740 | SNP | A→C | mutS | DNA mismatch repair protein MutS | L505R (CTG→CGG) | KT2440 GR18 ALE |
| 1,841,601 | DEL | Δ500 bp | [PP_RS08495],[PP_RS08500] | [PP_RS08495],[PP_RS08500] | | KT2440 GR18 TALE |
| 1,841,601 | DEL | Δ400 bp | [PP_RS08495] | [PP_RS08495] | | KT2440 GR18 TALE |
| 1,841,614 | DEL | Δ460 bp | [PP_RS08495],[PP_RS08500] | [PP_RS08495],[PP_RS08500] | | KT2440 GR18 TALE |
| 1,842,205 | SNP | G→C | PP_RS08500 | response regulator | H863Q (CAC→CAG) | KT2440 ALE |
| 1,842,521 | DEL | (T)5→4 | PP_RS08500 | response regulator | coding (2273/2754 nt) | KT2440 TALE |
| 1,842,601 | DEL | Δ36 bp | PP_RS08500 | response regulator | coding (2158-2193/2754 nt) | KT2440 TALE |
| 1,842,601 | DEL | Δ36 bp | PP_RS08500 | response regulator | coding (2158-2193/2754 nt) | KT2440 ALE |
| 1,842,632 | DEL | Δ11 bp | PP_RS08500 | response regulator | coding (2152-2162/2754 nt) | KT2440 GR18 ALE |
| 1,842,661 | DEL | Δ1 bp | PP_RS08500 | response regulator | coding (2133/2754 nt) | KT2440 ALE |
| 1,842,979 | SNP | G→C | PP_RS08500 | response regulator | Y605* (TAC→TAG) | KT2440 TALE |
| 1,843,067 | DEL | Δ4 bp | PP_RS08500 | response regulator | coding (1724-1727/2754 nt) | KT2440 GR18a ALE |
| 1,843,358 | SNP | A→G | PP_RS08500 | response regulator | L479P (CTG→CCG) | KT2440 TALE |
| 1,843,358 | SNP | A→T | PP_RS08500 | response regulator | L479Q (CTG→CAG) | KT2440 TALE |
| 1,843,367 | SNP | C→G | PP_RS08500 | response regulator | G476A (GGT→GCT) | KT2440 GR18 TALE |

| | | | | | | |
|---|---|---|---|---|---|---|
| 1,843,650 | SNP | C→T | PP_RS08500 | response regulator | E382K (GAA→AAA) | KT2440 GR18 ALE |
| 1,843,806 | SNP | A→G | PP_RS08500 | response regulator | S330P (TCT→CCT) | KT2440 GR18 ALE |
| 1,843,959 | SNP | C→T | PP_RS08500 | response regulator | A279T (GCA→ACA) | KT2440 GR18a ALE |
| 1,843,965 | INS | +A | PP_RS08500 | response regulator | coding (829/2754 nt) | KT2440 GR18 TALE |
| 1,844,100 | SNP | C→T | PP_RS08500 | response regulator | G232S (GGC→AGC) | KT2440 GR18a ALE |
| 1,844,131 | INS | +C | PP_RS08500 | response regulator | coding (663/2754 nt) | KT2440 GR18 TALE |
| 1,844,400 | INS | (AGGTCGC GGTGGTG GCC)1→2 | PP_RS08500 | response regulator | coding (394/2754 nt) | KT2440 GR18 ALE |
| 1,844,530 | DEL | Δ103 bp | PP_RS08500 | response regulator | coding (162-264/2754 nt) | KT2440 TALE |
| 1,844,624 | DEL | Δ56 bp | PP_RS08500 | response regulator | coding (115-170/2754 nt) | KT2440 GR18 TALE |
| 1,845,534 | SNP | A→G | PP_RS08505 | response regulator transcription factor | K183R (AAG→AGG) | KT2440 GR18 ALE |
| 1,846,310 | SNP | C→A | PP_RS08510 | sensor histidine kinase | R201S (CGC→AGC) | KT2440 GR18 TALE |
| 1,846,356 | SNP | T→G | PP_RS08510 | sensor histidine kinase | L216R (CTG→CGG) | KT2440 GR18 TALE |
| 1,846,397 | SNP | A→G | PP_RS08510 | sensor histidine kinase | T230A (ACG→GCG) | KT2440 GR18 TALE |
| 1,846,577 | SNP | G→C | PP_RS08510 | sensor histidine kinase | D290H (GAC→CAC) | KT2440 GR18 TALE |
| 1,846,895 | SNP | C→T | PP_RS08510 | sensor histidine kinase | P396S (CCG→TCG) | KT2440 GR18 TALE |
| 1,863,815 | SNP | C→T | PP_RS08585 | AI-2E family transporter | L141L (CTG→TTG) | KT2440 TALE |
| 1,863,815 | SNP | C→T | PP_RS08585 | AI-2E family transporter | L141L (CTG→TTG) | KT2440 ALE |
| 1,884,575 | INS | (A)9→10 | PP_RS08695, PP_RS08700 | hypothetical protein/SEL1-like repeat protein | intergenic (-908/-43) | KT2440 GR18 ALE |
| 1,970,753 | SNP | T→C | mtnA, gyrA | S-methyl-5-thioribose-1-phosphate isomerase/DNA gyrase subunit A | intergenic (+47/-319) | KT2440 GR18 ALE |

| | | | | | | |
|---|---|---|---|---|---|---|
| 1,989,645 | SNP | C→T | PP_RS09145 | glycosyltransferase | Q299* (CAA→TAA) | KT2440 TALE |
| 2,004,101 | DEL | Δ12,300 bp | [PP_RS09190],PP_RS09195 - PP_RS09230 | [PP_RS09190],PP_RS09195 - PP_RS09230 | P. putida KT2440 GR18 TALE | |
| 2,022,490 | SNP | C→T | gmd | GDP-mannose 4,6-dehydratase | R185C (CGC→TGC) | KT2440 ALE |
| 2,050,765 | SNP | G→T | PP_RS09380 | hypothetical protein | R80S (CGT→AGT) | KT2440 ALE |
| 2,082,258 | SNP | A→G | PP_RS09550, PP_RS09555 | MarR family transcriptional regulator/LysR family transcriptional regulator | intergenic (-37/+22) | KT2440 GR18 ALE |
| 2,101,751 | SNP | A→C | PP_RS09645 | hypothetical protein | L186R (CTG→CGG) | KT2440 GR18 ALE |
| 2,101,751 | SNP | A→C | PP_RS09645 | hypothetical protein | L186R (CTG→CGG) | KT2440 GR18 TALE |
| 2,168,080 | SNP | T→G | PP_RS09905 | hypothetical protein | W382G (TGG→GGG) | KT2440 GR18 TALE |
| 2,177,701 | AMP | 1,700 bp x 2 | PP_RS09965 | PP_RS09965 | duplication | KT2440 TALE |
| 2,177,701 | AMP | 1,700 bp x 2 | PP_RS09965 | PP_RS09965 | duplication | KT2440 ALE |
| 2,177,701 | AMP | 1,700 bp x 2 | PP_RS09965 | PP_RS09965 | duplication | KT2440 GR18 TALE |
| 2,182,594 | SNP | G→T | PP_RS09985, PP_RS09990 | hypothetical protein/hypothetical protein | intergenic (+12/-2) | KT2440 TALE |
| 2,213,079 | SNP | G→A | PP_RS10110 | cytochrome P450 | R404H (CGC→CAC) | KT2440 GR18 ALE |
| 2,372,151 | INS | (G)7→8 | PP_RS10800, PP_RS10805 | NAD-glutamate dehydrogenase/kinase/pyrophosphorylase | intergenic (+103/+9) | KT2440 GR18 ALE |
| 2,399,340 | SNP | C→T | PP_RS10905 | transporter substrate-binding domain-containing protein | S104N (AGC→AAC) | KT2440 GR18 ALE |
| 2,432,503 | SNP | A→G | PP_RS11065 | ATP-binding cassette domain-containing protein | E125E (GAA→GAG) | KT2440 GR18a ALE |
| 2,570,362 | SNP | G→A | PP_RS11700 | GntR family transcriptional regulator | N2N (AAC→AAT) | KT2440 GR18 ALE |
| 2,625,922 | DEL | Δ10 bp | PP_RS11995, PP_RS12000 | site-specific integrase/tRNA-His | intergenic (+100/-191) | KT2440 TALE |
| 2,625,922 | DEL | Δ10 bp | PP_RS11995, PP_RS12000 | site-specific integrase/tRNA-His | intergenic (+100/-191) | KT2440 ALE |
| 2,631,076 | SNP | G→T | lon | endopeptidase La | E181* (GAA→TAA) | KT2440 GR18 TALE |

| | | | | | | |
|---|---|---|---|---|---|---|
| 2,633,071 | DEL | Δ7 bp | lon, hupB | endopeptidase La/DNA-binding protein HU-beta | intergenic (+139/-8) | KT2440 GR18 TALE |
| 2,633,230 | SNP | C→T | hupB | DNA-binding protein HU-beta | T49I (ACC→ATC) | KT2440 GR18 TALE |
| 2,633,256 | SNP | C→T | hupB | DNA-binding protein HU-beta | R58C (CGT→TGT) | KT2440 GR18 TALE |
| 2,633,314 | SNP | C→T | hupB | DNA-binding protein HU-beta | P77L (CCA→CTA) | KT2440 GR18 TALE |
| 2,648,894 | SNP | G→T | PP_RS12110 | hypothetical protein | C72F (TGT→TTT) | KT2440 TALE |
| 2,699,014 | SNP | A→G | PP_RS12340 | helix-turn-helix transcriptional regulator | G66G (GGT→GGC) | KT2440 GR18 ALE |
| 2,752,455 | SNP | A→G | PP_RS12555 | efflux RND transporter periplasmic adaptor subunit | P4P (CCA→CCG) | KT2440 GR18 ALE |
| 2,831,084 | SNP | A→C | PP_RS12965, PP_RS12970 | nucleotidyltransferase family protein/ArsR family transcriptional regulator | intergenic (-142/+141) | KT2440 TALE |
| 2,831,084 | SNP | A→C | PP_RS12965, PP_RS12970 | nucleotidyltransferase family protein/ArsR family transcriptional regulator | intergenic (-142/+141) | KT2440 ALE |
| 2,864,458 | SNP | C→T | PP_RS13160 | OprD family porin | A309T (GCC→ACC) | KT2440 GR18 ALE |
| 2,974,486 | DEL | Δ2 bp | PP_RS13565 | IclR family transcriptional regulator | coding (419-420/777 nt) | KT2440 ALE |
| 2,975,957 | INS | (G)6→7 | PP_RS13570 | Gfo/Idh/MocA family oxidoreductase | coding (975/1053 nt) | KT2440 GR18 ALE |
| 3,045,232 | SNP | C→A | pstC | phosphate ABC transporter permease subunit PstC | A258A (GCC→GCA) | KT2440 GR18a ALE |
| 3,194,798 | SNP | C→T | PP_RS14565 | APC family permease | L24L (CTG→CTA) | KT2440 GR18 ALE |
| 3,366,401 | AMP | 17,200 bp x 2 | PP_RS15365 - PP_RS28550,[PP_RS15460] | PP_RS15365 - PP_RS28550,[PP_RS15460] | duplication | KT2440 ALE |
| 3,366,501 | AMP | 17,100 bp x 2 | PP_RS15365 - PP_RS28550,[PP_RS15460] | PP_RS15365 - PP_RS28550,[PP_RS15460] | duplication | KT2440 TALE |
| 3,366,501 | AMP | 6,200 bp x 2 | PP_RS15365 - PP_RS15390,[PP_RS15395] | PP_RS15365 - PP_RS15390,[PP_RS15395] | duplication | KT2440 ALE |
| 3,366,501 | AMP | 17,000 bp x 2 | PP_RS15365 - PP_RS28550 | PP_RS15365 - PP_RS28550 | duplication | KT2440 ALE |
| 3,366,601 | AMP | 17,000 bp x 2 | PP_RS15365 - PP_RS28550,[PP_RS15460] | PP_RS15365 - PP_RS28550,[PP_RS15460] | duplication | KT2440 ALE |
| 3,366,601 | AMP | 17,000 bp x 2 | PP_RS15365 - PP_RS28550,[PP_RS15460] | PP_RS15365 - PP_RS28550,[PP_RS15460] | duplication | KT2440 GR18 ALE |

| | | | | | | |
|---|---|---|---|---|---|---|
| 3,367,801 | AMP | 15,800 bp x 2 | [PP_RS15365],PP_RS15370 - PP_RS28550,[PP_RS15460] | [PP_RS15365],PP_RS15370 - PP_RS28550,[PP_RS15460] | duplication | KT2440 TALE |
| 3,373,101 | AMP | 10,500 bp x 2 | [PP_RS15400],PP_RS15405 - PP_RS28550,[PP_RS15460] | [PP_RS15400],PP_RS15405 - PP_RS28550,[PP_RS15460] | duplication | KT2440 ALE |
| 3,396,064 | SNP | G→A | PP_RS15530 | MaoC family dehydratase | E76K (GAG→AAG) | KT2440 GR18 ALE |
| 3,447,286 | SNP | A→C | PP_RS15865 | lysis system i-spanin subunit Rz | S18R (AGC→CGC) | KT2440 GR18a ALE |
| 3,613,139 | SNP | G→A | PP_RS16600 | Ig-like domain-containing protein | P182L (CCG→CTG) | KT2440 GR18 ALE |
| 3,631,655 | SNP | G→A | PP_RS16685 | DUF1329 domain-containing protein | A183A (GCG→GCA) | KT2440 GR18 ALE |
| 3,662,938 | SNP | G→T | PP_RS16815, PP_RS16820 | acyl-CoA/acyl-ACP dehydrogenase/LysR family transcriptional regulator | intergenic (-114/+81) | KT2440 GR18 ALE |
| 3,697,578 | SNP | T→C | PP_RS17020 | lysylphosphatidylglycerol synthase domain-containing protein | Y298C (TAC→TGC) | KT2440 GR18 ALE |
| 3,749,162 | SNP | G→A | PP_RS17270 | Hsp20/alpha crystallin family protein | R99R (CGC→CGT) | KT2440 TALE |
| 3,763,200 | SNP | G→A | PP_RS17330 | DUF4198 domain-containing protein | S148N (AGC→AAC) | KT2440 GR18 ALE |
| 3,822,062 | SNP | G→A | PP_RS17590 | MFS transporter | G374G (GGC→GGT) | KT2440 GR18 ALE |
| 3,828,301 | AMP | 245,600 bp x 2 | PP_RS17615 - PP_RS18625 | PP_RS17615 - PP_RS18625 | duplication | KT2440 GR18a ALE |
| 3,828,301 | AMP | 143,500 bp x 2 | PP_RS17615 - PP_RS18205 | PP_RS17615 - PP_RS18205 | duplication | KT2440 GR18a ALE |
| 3,828,301 | AMP | 245,100 bp x 2 | PP_RS17615 - _ PP_RS18620,[PP_RS18625] | PP_RS17615 - _ PP_RS18620,[PP_RS18625] | duplication | KT2440 GR18a ALE |
| 3,828,301 | AMP | 245,200 bp x 2 | PP_RS17615 - PP_RS18625 | PP_RS17615 - PP_RS18625 | duplication | KT2440 GR18a ALE |
| 3,951,179 | SNP | T→C | PP_RS18130, PP_RS18135 | ABC transporter substrate-binding protein/SCO family protein | intergenic (-143/+52) | KT2440 GR18 ALE |
| 3,973,001 | AMP | 100,800 bp x 2 | PP_RS18215 - PP_RS18625 | PP_RS18215 - PP_RS18625 | duplication | KT2440 GR18a ALE |
| 4,013,344 | DEL | (T)7→6 | PP_RS18400, PP_RS18405 | hydroxymethylglutaryl-CoA lyase/MgtC/SapB family protein | intergenic (+280/+253) | KT2440 GR18 TALE |
| 4,013,466 | DEL | (A)7→6 | PP_RS18400, PP_RS18405 | hydroxymethylglutaryl-CoA lyase/MgtC/SapB family protein | intergenic (+402/+131) | KT2440 TALE |

| | | | | | | |
|---|---|---|---|---|---|---|
| 4,028,211 | SNP | T→C | PP_RS18460 | PAS domain S-box protein | E73E (GAA→GAG) | KT2440 GR18 ALE |
| 4,047,034 | SNP | T→C | PP_RS18540 | hypothetical protein | R66R (CGT→CGC) | KT2440 GR18 ALE |
| 4,063,990 | SNP | C→T | PP_RS18605 | bifunctional diguanylate cyclase/phosphodiesterase | G117S (GGT→AGT) | KT2440 GR18 ALE |
| 4,067,105 | INS | (C)6→7 | PP_RS18615 | efflux RND transporter permease subunit | coding (1959/3108 nt) | KT2440 GR18 ALE |
| 4,140,900 | SNP | G→A | PP_RS18925 | amino acid synthesis family protein | L176L (CTG→CTA) | KT2440 GR18 ALE |
| 4,157,832 | SNP | C→T | PP_RS19005 | LysR family transcriptional regulator | G16D (GGC→GAC) | KT2440 GR18 ALE |
| 4,179,035 | SNP | T→C | PP_RS19085 | hypothetical protein | Y536Y (TAT→TAC) | KT2440 ALE |
| 4,183,728 | INS | (G)5→6 | PP_RS19105 | AAA family ATPase | coding (1434/1932 nt) | KT2440 GR18 ALE |
| 4,187,531 | SNP | T→C | PP_RS19120 | hypothetical protein | L274P (CTG→CCG) | KT2440 GR18 ALE |
| 4,204,431 | SNP | G→A | PP_RS19210 | DUF3320 domain-containing protein | I1115I (ATC→ATT) | KT2440 GR18 ALE |
| 4,233,513 | INS | (GAGCAG)7 →8 | PP_RS19320, PP_RS19325 | helix-turn-helix transcriptional regulator/diguanylate cyclase | intergenic (+190/+46) | KT2440 TALE |
| 4,233,513 | INS | (GAGCAG)7 →8 | PP_RS19320, PP_RS19325 | helix-turn-helix transcriptional regulator/diguanylate cyclase | intergenic (+190/+46) | KT2440 ALE |
| 4,307,298 | SNP | T→C | PP_RS19670 | LysR family transcriptional regulator | S235G (AGC→GGC) | KT2440 GR18 ALE |
| 4,341,274 | SNP | C→A | PP_RS19850 | polyamine ABC transporter substrate-binding protein | T310N (ACT→AAT) | KT2440 TALE |
| 4,341,274 | SNP | C→A | PP_RS19850 | polyamine ABC transporter substrate-binding protein | T310N (ACT→AAT) | KT2440 ALE |
| 4,348,705 | SNP | A→G | ltrA, galU | group II intron reverse transcriptase/maturase/UTP--glucose-1-phosphate uridylyltransferase GalU | intergenic (-70/+404) | KT2440 TALE |
| 4,348,705 | SNP | A→G | ltrA, galU | group II intron reverse transcriptase/maturase/UTP--glucose-1-phosphate uridylyltransferase GalU | intergenic (-70/+404) | KT2440 ALE |

| | | | | | | |
|---|---|---|---|---|---|---|
| 4,348,705 | SNP | A→G | ltrA, galU | group II intron reverse transcriptase/maturase/UTP--glucose-1-phosphate uridylyltransferase GalU | intergenic (-70/+404) | KT2440 GR18 ALE |
| 4,348,705 | SNP | A→G | ltrA, galU | group II intron reverse transcriptase/maturase/UTP--glucose-1-phosphate uridylyltransferase GalU | intergenic (-70/+404) | KT2440 GR18 TALE |
| 4,348,705 | SNP | A→G | ltrA, galU | group II intron reverse transcriptase/maturase/UTP--glucose-1-phosphate uridylyltransferase GalU | intergenic (-70/+404) | KT2440 GR18a ALE |
| 4,380,648 | SNP | G→A | PP_RS20075 | hypothetical protein | T714T (ACC→ACT) | KT2440 GR18 TALE |
| 4,470,054 | SNP | G→A | PP_RS20600 | MFS transporter | A379V (GCA→GTA) | KT2440 GR18 ALE |
| 4,536,201 | AMP | 2,000 bp x 2 | [PP_RS20920],tnpB,PP_RS20930 | [PP_RS20920],tnpB,PP_RS20930 | duplication | KT2440 GR18 TALE |
| 4,571,846 | INS | (C)8→9 | PP_RS21060 | malto-oligosyltrehalose synthase | coding (1710/2775 nt) | KT2440 GR18 ALE |
| 4,583,581 | DEL | (G)11→10 | treS, PP_RS21095 | maltose alpha-D-glucosyltransferase/alpha-1,4-glucan--maltose-1-phosphate maltosyltransferase | intergenic (-154/+17) | KT2440 GR18 ALE |
| 4,597,535 | SNP | T→C | tssM | type VI secretion system membrane subunit TssM | E758E (GAA→GAG) | KT2440 GR18 ALE |
| 4,612,826 | DEL | (G)6→5 | vgrG | type VI secretion system tip protein VgrG | pseudogene (1714/1985 nt) | KT2440 GR18 ALE |
| 4,628,925 | SNP | T→C | PP_RS21265 | hypothetical protein | F85F (TTT→TTC) | KT2440 GR18 TALE |
| 4,635,167 | SNP | C→T | uvrY | UvrY/SirA/GacA family response regulator transcription factor | E197K (GAA→AAA) | KT2440 GR18 TALE |
| 4,635,282 | SNP | C→T | uvrY | UvrY/SirA/GacA family response regulator transcription factor | M158I (ATG→ATA) | KT2440 TALE |
| 4,635,333 | DEL | Δ1 bp | uvrY | UvrY/SirA/GacA family response regulator transcription factor | coding (423/639 nt) | KT2440 ALE |
| 4,635,341 | SNP | G→A | uvrY | UvrY/SirA/GacA family response regulator transcription factor | Q139* (CAG→TAG) | KT2440 TALE |
| 4,635,365 | SNP | G→A | uvrY | UvrY/SirA/GacA family response regulator transcription factor | Q131* (CAG→TAG) | KT2440 TALE |

| | | | | | | |
|---|---|---|---|---|---|---|
| 4,635,487 | SNP | G→A | uvrY | UvrY/SirA/GacA family response regulator transcription factor | P90L (CCC→CTC) | KT2440 GR18 ALE |
| 4,635,516 | DEL | Δ6 bp | uvrY | UvrY/SirA/GacA family response regulator transcription factor | coding (235-240/639 nt) | KT2440 GR18a ALE |
| 4,635,583 | SNP | G→A | uvrY | UvrY/SirA/GacA family response regulator transcription factor | P58L (CCC→CTC) | KT2440 ALE |
| 4,635,745 | SNP | A→C | uvrY | UvrY/SirA/GacA family response regulator transcription factor | V4G (GTC→GGC) | KT2440 TALE |
| 4,635,745 | SNP | A→C | uvrY | UvrY/SirA/GacA family response regulator transcription factor | V4G (GTC→GGC) | KT2440 ALE |
| 4,635,745 | SNP | A→C | uvrY | UvrY/SirA/GacA family response regulator transcription factor | V4G (GTC→GGC) | KT2440 GR18 ALE |
| 4,635,765 | SNP | T→G | uvrY, PP_RS21315 | UvrY/SirA/GacA family response regulator transcription factor/helix-turn-helix domain-containing protein | intergenic (-10/+54) | KT2440 TALE |
| 4,635,766 | SNP | C→G | uvrY, PP_RS21315 | UvrY/SirA/GacA family response regulator transcription factor/helix-turn-helix domain-containing protein | intergenic (-11/+53) | KT2440 GR18 TALE |
| 4,656,320 | SNP | G→A | PP_RS21405 | NADH-quinone oxidoreductase subunit A | E46K (GAA→AAA) | KT2440 GR18 ALE |
| 4,716,608 | SNP | T→C | PP_RS21670 | ATP-binding protein | D613D (GAT→GAC) | KT2440 GR18 ALE |
| 4,739,196 | SNP | G→C | sdhC, gltA | succinate dehydrogenase, cytochrome b556 subunit/citrate synthase | intergenic (-75/-278) | KT2440 ALE |
| 4,831,403 | SNP | C→T | PP_RS22040, PP_RS22045 | non-ribosomal peptide synthetase/RNA polymerase factor sigma-70 | intergenic (-214/-92) | KT2440 GR18a ALE |
| 4,888,387 | SNP | G→A | gcl | glyoxylate carboligase | A482A (GCG→GCA) | KT2440 GR18 ALE |
| 4,897,623 | INS | (C)7→8 | PP_RS22360 | class I SAM-dependent methyltransferase | coding (921/957 nt) | KT2440 GR18 ALE |
| 4,963,501 | DEL | Δ1,000 bp | [PP_RS22695],[fleQ] | [PP_RS22695],[fleQ] | | KT2440 TALE |
| 4,964,047 | DEL | Δ4 bp | fleQ | transcriptional regulator FleQ | coding (1308-1311/1476 nt) | KT2440 TALE |
| 4,964,207 | SNP | G→T | fleQ | transcriptional regulator FleQ | S384* (TCG→TAG) | KT2440 TALE |
| 4,964,267 | DEL | Δ22 bp | fleQ | transcriptional regulator FleQ | coding (1070-1091/1476 nt) | KT2440 ALE |

| | | | | | | |
|---|---|---|---|---|---|---|
| 4,964,322 | DEL | Δ1 bp | fleQ | transcriptional regulator FleQ | coding (1036/1476 nt) | KT2440 ALE |
| 4,964,501 | SNP | G→A | fleQ | transcriptional regulator FleQ | T286M (ACG→ATG) | KT2440 ALE |
| 4,964,622 | SNP | C→T | fleQ | transcriptional regulator FleQ | E246K (GAA→AAA) | KT2440 ALE |
| 4,964,661 | SNP | G→A | fleQ | transcriptional regulator FleQ | R233C (CGT→TGT) | KT2440 ALE |
| 4,964,733 | SNP | C→A | fleQ | transcriptional regulator FleQ | E209* (GAG→TAG) | KT2440 TALE |
| 4,964,820 | INS | +A | fleQ | transcriptional regulator FleQ | coding (538/1476 nt) | KT2440 TALE |
| 4,964,916 | SNP | C→T | fleQ | transcriptional regulator FleQ | G148S (GGC→AGC) | KT2440 TALE |
| 5,010,127 | SNP | T→C | PP_RS22900, PP_RS22905 | hypothetical protein/tyrosine-type recombinase/integrase | intergenic (+487/-364) | KT2440 GR18 ALE |
| 5,070,922 | SNP | T→C | PP_RS23205, PP_RS23210 | TauD/TfdA family dioxygenase/LysR family transcriptional regulator | intergenic (-36/-210) | KT2440 GR18 ALE |
| 5,073,252 | SNP | C→T | PP_RS23220 | hypothetical protein | A122T (GCC→ACC) | KT2440 GR18 ALE |
| 5,080,226 | SNP | G→A | csrA, PP_RS23295 | carbon storage regulator CsrA/aspartate kinase | intergenic (-91/+77) | KT2440 GR18 ALE |
| 5,097,817 | SNP | G→C | acs | acetate--CoA ligase | D482E (GAC→GAG) | KT2440 TALE |
| 5,172,783 | INS | (C)5→6 | fadD1 | long-chain-fatty-acid--CoA ligase FadD1 | coding (177/1698 nt) | KT2440 GR18 ALE |
| 5,300,103 | SNP | G→A | recB | exodeoxyribonuclease V subunit beta | P1224S (CCA→TCA) | KT2440 TALE |
| 5,300,103 | SNP | G→A | recB | exodeoxyribonuclease V subunit beta | P1224S (CCA→TCA) | KT2440 ALE |
| 5,317,968 | SNP | G→A | PP_RS24400 | acetolactate synthase 3 large subunit | P469L (CCG→CTG) | KT2440 GR18 ALE |
| 5,322,212 | SNP | T→C | mrcB | penicillin-binding protein 1B | D347G (GAT→GGT) | KT2440 GR18 ALE |
| 5,338,322 | DEL | Δ30 bp | PP_RS24485, PP_RS24490 | sigma-54 dependent transcriptional regulator/polynucleotide adenylyltransferase PcnB | intergenic (+329/-422) | KT2440 TALE |

| | | | | | | |
|---|---|---|---|---|---|---|
| 5,399,567 | DEL | (T)5→4 | PP_RS24755 | AAA family ATPase | coding (380/1437 nt) | KT2440 TALE |
| 5,399,567 | DEL | (T)5→4 | PP_RS24755 | AAA family ATPase | coding (380/1437 nt) | KT2440 ALE |
| 5,476,595 | DEL | (GCT)3→2 | PP_RS25115 | bifunctional DedA family/phosphatase PAP2 family protein | coding (825-827/1317 nt) | KT2440 TALE |
| 5,549,195 | DEL | (GAATTTTTT)2→1 | rnr, PP_RS25470 | ribonuclease R/tRNA-Leu | intergenic (-142/-123) | KT2440 GR18 TALE |
| 5,643,470 | SNP | T→C | PP_RS25850 | NAD(P)/FAD-dependent oxidoreductase | S45P (TCC→CCC) | KT2440 GR18 ALE |
| 5,817,498 | SNP | G→A | PP_RS26545 | PilT/PilU family type 4a pilus ATPase | N148N (AAC→AAT) | KT2440 GR18 ALE |
| 5,821,926 | SNP | T→C | metW | methionine biosynthesis protein MetW | W177R (TGG→CGG) | KT2440 ALE |
| 5,842,840 | SNP | T→C | PP_RS26680 | coniferyl aldehyde dehydrogenase | V470A (GTC→GCC) | KT2440 GR18 ALE |
| 5,874,795 | SNP | C→G | ilvA | threonine ammonia-lyase, biosynthetic | G459A (GGC→GCC) | KT2440 GR18 TALE |
| 5,874,805 | SNP | G→A | ilvA | threonine ammonia-lyase, biosynthetic | R456C (CGC→TGC) | KT2440 TALE |
| 5,891,891 | SNP | T→C | PP_RS26915 | sigma 54-interacting transcriptional regulator | T245A (ACC→GCC) | KT2440 GR18 ALE |
| 5,998,486 | SNP | A→C | PP_RS27385, PP_RS27390 | hypothetical protein/hydrolase | intergenic (-88/+30) | KT2440 TALE |
| 6,040,002 | SNP | A→T | PP_RS27575 | exodeoxyribonuclease III | I3F (ATC→TTC) | KT2440 GR18 TALE |
| 6,040,244 | SNP | G→A | PP_RS27575 | exodeoxyribonuclease III | L83L (CTG→CTA) | KT2440 TALE |
| 6,040,321 | DEL | (TGC)3→2 | PP_RS27575 | exodeoxyribonuclease III | coding (326-328/780 nt) | KT2440 GR18 TALE |
| 6,040,668 | SNP | A→C | PP_RS27575 | exodeoxyribonuclease III | T225P (ACC→CCC) | KT2440 GR18 TALE |
| 6,040,772 | DEL | Δ1 bp | PP_RS27575 | exodeoxyribonuclease III | coding (777/780 nt) | KT2440 TALE |
| 6,077,816 | DEL | Δ98 bp | PP_RS27755, PP_RS27760 | phosphate ABC transporter substrate-binding protein PstS/MFS transporter | intergenic (-118/+23) | KT2440 ALE |
| 6,116,136 | SNP | C→T | lpdA | dihydrolipoyl dehydrogenase | G115G (GGC→GGT) | KT2440 GR18 ALE |

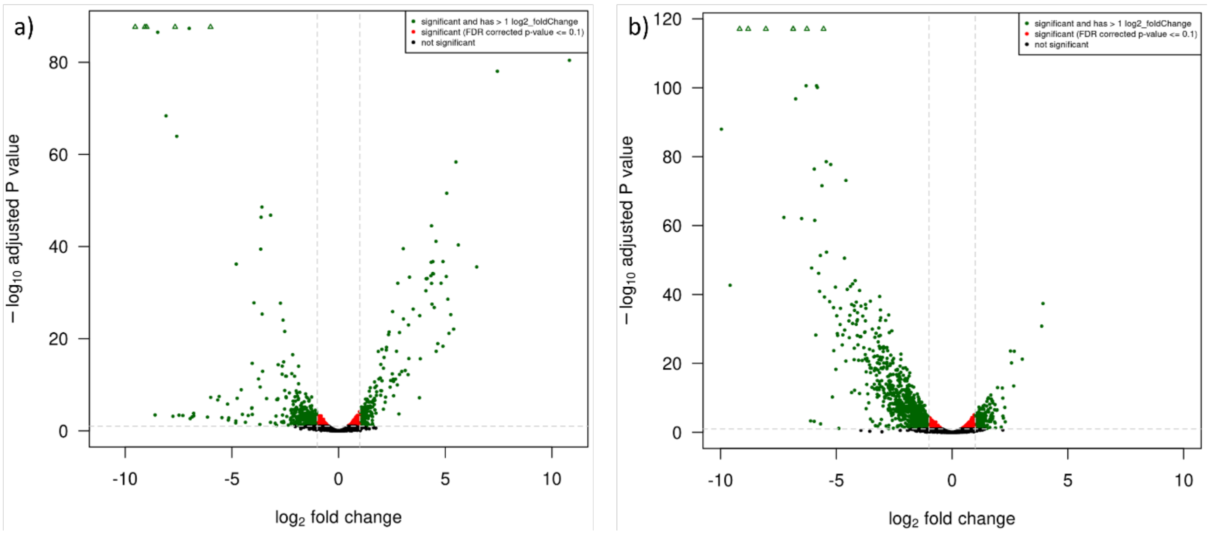


**Figure S 3: Volcano plot showing differentially expressed genes a) during growth of *P. putida* KT2440 on acetate vs. on glucose and b) during growth of *P. putida* KT2440 vs. *P. putida* KT2440 TALE during growth on acetate.**
